# Microbial oral-gut translocation in advanced chronic liver disease is linked to exacerbation of intestinal barrier dysfunction and hepatic fibrosis

**DOI:** 10.1101/2025.09.23.677032

**Authors:** Shen Jin, Aurelie Cenier, Daniela Wetzel, Bethlehem Arefaine, Mar Moreno-Gonzalez, Marilena Stamouli, Merianne Mohamad, Mariia Lupatsii, Emilio Ríos, Sunjae Lee, Ane Zamalloa, Shilpa Chokshi, Adil Mardinoglu, Saeed Shoaie, Naiara Beraza, Vishal C Patel, Melanie Schirmer

## Abstract

While microbiome perturbations are associated with advanced chronic liver disease (ACLD), microbial disease mechanisms are poorly understood. Using multi-omics analyses of paired saliva and faecal samples from an ACLD cohort, we identified next-to-identical oral and gut bacterial strains (including *Veillonella* and *Streptococcus spp.*) which increased in absolute abundance in the gut of ACLD patients. These translocators uniquely encoded a collagenase-like proteinase (*prtC*) with the potential for gut barrier disruption and *prtC* faecal abundance was a robust ACLD biomarker (auPR=0.91). CCl_4_-treated mice inoculated with *Veillonella* and *Streptococcus prtC-*encoding patient isolates showed exacerbation of gut barrier impairment and hepatic fibrosis. Furthermore, faecal collagenase activity was increased in ACLD patients and experimentally confirmed for the *prtC* gene from translocating *Veillonella parvula*. Overall, our study establishes mechanistic links between oral-gut translocation and ACLD pathobiology, and identifies the oral microbiome as an important contributing factor with potential for microbial diagnostics and therapeutics.

## Introduction

Advanced chronic liver disease (ACLD) occurs as a consequence of the histological development of cirrhosis and is a global disease burden [1] accounting for over two million deaths every year [2]. It results from chronic liver inflammation, where normal hepatic parenchyma is replaced by scar tissue and regenerative nodules [3, 4], and comprises several stages. The first stage is compensated ACLD (cACLD), where hepatic synthetic function is maintained and patients are mostly asymptomatic. Subsequently, acutely decompensated ACLD (AD) is driven by worsening portal hypertension and portosystemic shunting, and characterised by synthetic failure and the acute and often unpredictable onset of complications including ascites, jaundice, hepatic encephalopathy (HE), variceal haemorrhage, heightened risk of infection and hepatocellular carcinoma [5]. The most severe form is acute-on-chronic liver failure (ACLF), which occurs in both cACLD and AD patients and is marked by systemic inflammation, (multi-)organ failure, and high short-term mortality (30-days mortality rate in >57% of patients) [6, 7]. Current standard of care mainly targets complications, while interventions to prevent disease progression [8] and new approaches for early diagnosis of asymptomatic patients with advanced hepatic fibrosis are lacking [9, 10].

Several studies have identified gut microbial perturbations associated with ACLD pathobiology. Gut microbiome diversity is decreased and characterised by a loss of bacterial commensals, such as *Lachnospiraceae*, *Bacteroidaceae* and *Ruminococcaceae*, while the relative abundance of opportunistic pathogens increases, including *Enterobacteriaceae*, *Enterococcaceae*, *Veillonellaceae*, and *Streptococcaceae* [11–17] [18]. Microbial factors can also predict ACLD [19], where the abundances of 19 species [20] was used to distinguish between cirrhosis patients and healthy controls (auROC: 0.86). Furthermore, antibiotic resistance genes [21] and microbially produced aromatic and branched-chain amino acids [20] are associated with ACLD, while increased microbial fatty-acid biosynthesis and glycan degradation activity is associated with cirrhosis severity and hepatic damage [22]. While this implicates microbiome perturbations in disease, mechanistic insights into how this contributes to ACLD pathobiology are still missing.

Recently, a pronounced increase of bacteria typical for the oral cavities was identified, where *Veillonella*, *Streptococcus*, and *Prevotella spp.* were found in high relative abundance in ACLD patients’ faeces [12, 23]. This was also reported for other diseases, including ulcerative colitis [24], colorectal cancer [25, 26], rheumatoid arthritis, type-1 diabetes [27] [28] and hypertension [29], suggesting that microbiome oral-gut translocation may play an important role in these disease states [30] (reviewed in [31]). Rifaximin-α, a gut-targeted therapy to prevent HE in AD, reduces oralisation of the gut microbiome and suppresses bacteria with intestinal mucus degradation capabilities, which may ameliorate HE symptoms by promoting gut barrier integrity [32]. Another bacterium typical for the oral cavity, *Veillonella parvula,* degrades immunosuppressive thiopurine drugs through *xdhA* xanthine dehydrogenase potentially impacting therapeutic efficacy in inflammatory bowel disease (IBD) patients [24]. However, it is still unclear if these bacteria are actually translocating from the mouth and colonise the gut during disease or if this increase in oral bacterial abundance is due to a decrease in total gut bacterial load [33]. Paired saliva and faecal samples with metagenomic strain profiling are essential to investigate the oral origin of these atypical gut bacteria, but these data are currently lacking for the majority of cohort studies.

Here, we investigated the role of oral-gut translocation in ACLD using metagenomic data from paired saliva and faecal samples. Using reference- and assembly-based analyses, we computationally identified co-occurring bacterial strains in saliva and faeces, obtained viable next-to-identical clinical isolates and experimentally confirmed increases in absolute abundance of oral microbes in the ACLD faecal samples. By integrating microbiome strain and functional profiles with detailed clinical data, including disease severity indices (Child-Pugh and Model for End-Stage Liver Disease [MELD] scores) and a small intestinal epithelial cell damage marker (plasma Fatty Acid Binding Protein 2 [FABP2]), we identified a bacterial collagenase-like proteinase (*prtC*) associated with disease pathobiology. This *prtC* gene was uniquely shared by oral-gut translocating bacteria and its stool abundance emerged as a robust biomarker of ACLD (auPR=0.91, auROC=0.89; external validation cohort: auPR=0.93, auROC=0.93). We further investigated the role of these translocators using a preclinical mouse model of hepatic fibrosis. Gavaging oral patient isolates of *Veillonella* and *Streptococcus spp.* into CCl_4_-treated mice exacerbated gut barrier impairment (including increases in colonic albumin levels and mislocalization of E-cadherin and occludin), which was accompanied by increased hepatic and intestinal fibrosis (measured by Sirius red staining of liver tissue) and small intestinal bacterial overgrowth. Collagenase activity was also increased in ACLD patients’ faecal supernatant compared to healthy controls. Furthermore, *in vitro* expression of the *Veillonella parvula prtC* in *Escherichia coli* confirmed collagenase activity of the implicated strains. Collectively, our study links microbial strains and genes involved in oral-gut translocation to ACLD pathobiology.

## Results

### Oral microbiome shifts are detected at early disease stages and oral-gut similarity increases with ACLD progression

Saliva and faecal samples from 86 ACLD patients and two control groups consisting of 52 healthy participants and 14 septic patients with no underlying cirrhosis were analysed (**Figure 1a**, see **Table 1** for clinical characteristics). Sepsis patients formed our second control group as they showed similar characteristics in hospitalisation, age, antibiotic usage and PPI exposure compared to ACLD patients, while representing a distinct disease with different disease mechanisms. ACLD patients were grouped based on disease severity, including patients with cACLD, AD and ACLF, and metagenomic sequencing data from 266 samples (n_faecal_=143, n_saliva_=123) was used to identify taxonomic and functional microbiome signals associated with oral-gut translocation.

**Figure 1:**
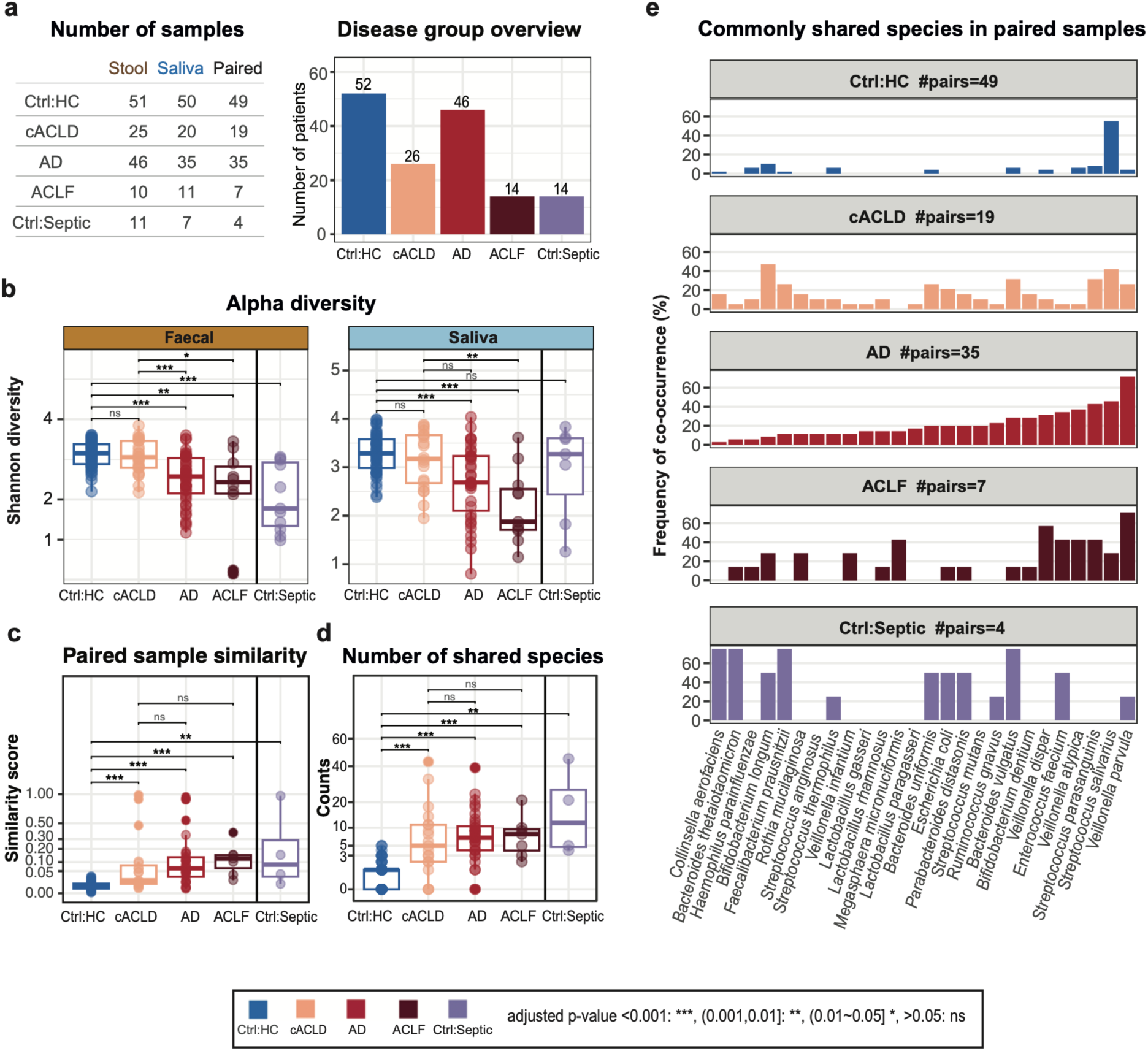
Oral-gut translocation increases as ACLD severity worsens. **(a)** Cohort overview. Number of faecal, saliva and paired samples and number of patients per disease group. ACLD patients were stratified according to disease stage ranging from compensated ACLD (cACLD) and acutely decompensated ACLD (AD) to acute-on-chronic liver failure (ACLF). Two control groups were included: healthy patients (Ctrl:HC) and septic patients (Ctrl:Septic). **(b)** Alpha diversity (Shannon index) decreased in faeces and saliva as ACLD severity increased (Wilcoxon, ns = not significant, p-value adjusted with Benjamini Hochberg [BH]). **(c)** Similarity (1 - Bray-Curtis dissimilarity) between paired faecal and saliva samples stratified by disease group (Wilcoxon, ns = not significant, p-value adjusted with BH) **(d)** Number of shared species in paired faecal and saliva samples increased significantly as disease severity increased (Wilcoxon, based on reference-based species profiles, p-value adjusted with BH). **(e)** Identification of microbial species commonly shared between saliva and faecal samples. Y-axis indicates percentage of samples where the species was detected in both paired sample types. Each column represents a species ordered by their frequency in the AD group.

**Table 1:** Summary of patient clinical characteristics. Normally distributed values are denoted with (*) and are presented as mean ± standard deviation (SD); non-normally distributed values are presented as median (interquartile range).

While gut microbiome diversity was decreased in both ACLD and septic patients, decreases in oral microbiome diversity were only observed in ACLD patients and not in healthy controls or sepsis patients (**Figure 1b**). AD and ACLF patients with more severe disease showed the most drastic decrease in alpha diversity of both the oral and gut microbiome. Concurrently, microbial dysbiosis as measured by the dysbiosis index [34] increased in both faeces and saliva in all ACLD groups **(Figure S1a)**. Importantly, oral microbial dysbiosis seemed to occur at earlier disease severity stages with significant changes already occurring in cACLD patients compared to Ctrl:HC (**Figure S1a,** Wilcoxon p=0.0012). While the dysbiosis score reflects the dissimilarity of a given sample from the average healthy microbiome, the dysbiosis score distribution *within* healthy individuals reflects the inter-individual variations among this population for a given body site. Interestingly, dysbiosis scores were generally lower for healthy oral microbiomes compared to healthy gut microbiomes (median_saliva_=0.55, median_faeces_=0.69, **Figure S1a**), suggesting that the oral microbiome across healthy individuals is more similar than their gut microbiomes.

We next compared intra-patient oral-gut similarity using paired saliva and faecal samples. Microbiome similarity between paired saliva and faecal samples was significantly increased in all ACLD groups (**Figure 1c**, 1 - Bray-Curtis dissimilarity, Wilcoxon, adjusted p-value <0.001, Benjamini-Hochberg (BH) corrected). The number of shared species also increased in the ACLD groups (**Figure 1d**, reference-based profiling), with *Veillonella parvula, Veillonella atypica*, *Streptococcus salivarius* and *Streptococcus parasanguinis* being the most commonly shared species in paired saliva and faecal samples **(Figure 1e)**. Importantly, this increase in shared species was independent of the number of patients in each disease group (**Figure S1b**). *Veillonella* can ectopically colonise the gut during inflammation through nitrate respiration using the narGHJI operon [35] and host-derived nitrate boosts *Escherichia coli* growth [36]. We detected the nar operon in 8 assembled Metagenomic Species Pangenomes (MSPs), including *E. coli*, *V. parvula* and *V. dispar* (**Figure S1c**), which were among the most commonly shared oral-gut species in ACLD. Overall, this implicates the oral microbiome in ACLD progression, where oral dysbiosis may precede changes in the gut microbiome.

### Co-occurring strains are detected in paired saliva and faecal samples in ACLD

We next performed strain-level profiling to further support the presence of oral-gut translocation associated with ACLD. As the gastro-intestinal tract provides a natural route for oral microbes to reach the gut, we focused our analysis on translocation from the mouth to the gut. First, we identified common members of the saliva microbiome (present in >20% of saliva samples) among the shared species (n_shared_=26, **Figure 1e, Table S1**), resulting in 9 candidate translocators (**Figure S1d**). Two strategies were used to evaluate strain similarity: (1) phylogenetic distance between strains based on single nucleotide polymorphisms (SNP) in marker genes [37] [38] and (2) similarity of gene presence/absence patterns based on pangenome analysis [37, 39]. These complementary strategies balance sequencing coverage requirements for SNP analysis with reference genome requirements for pangenome analysis. For all candidate translocator species except *Veillonella infantium*, at least one type of strain profile was available (**Figure S1d**). For the remaining eight translocator species our analyses showed that within the same individual (i.e. in paired samples) oral and gut strains were highly similar, while oral and gut strains in samples from different individuals (i.e. unpaired samples) showed significantly greater strain diversity (**Figures 2a**+**S1e**), providing computational evidence for bacterial oral-gut translocation.

**Figure 2:**
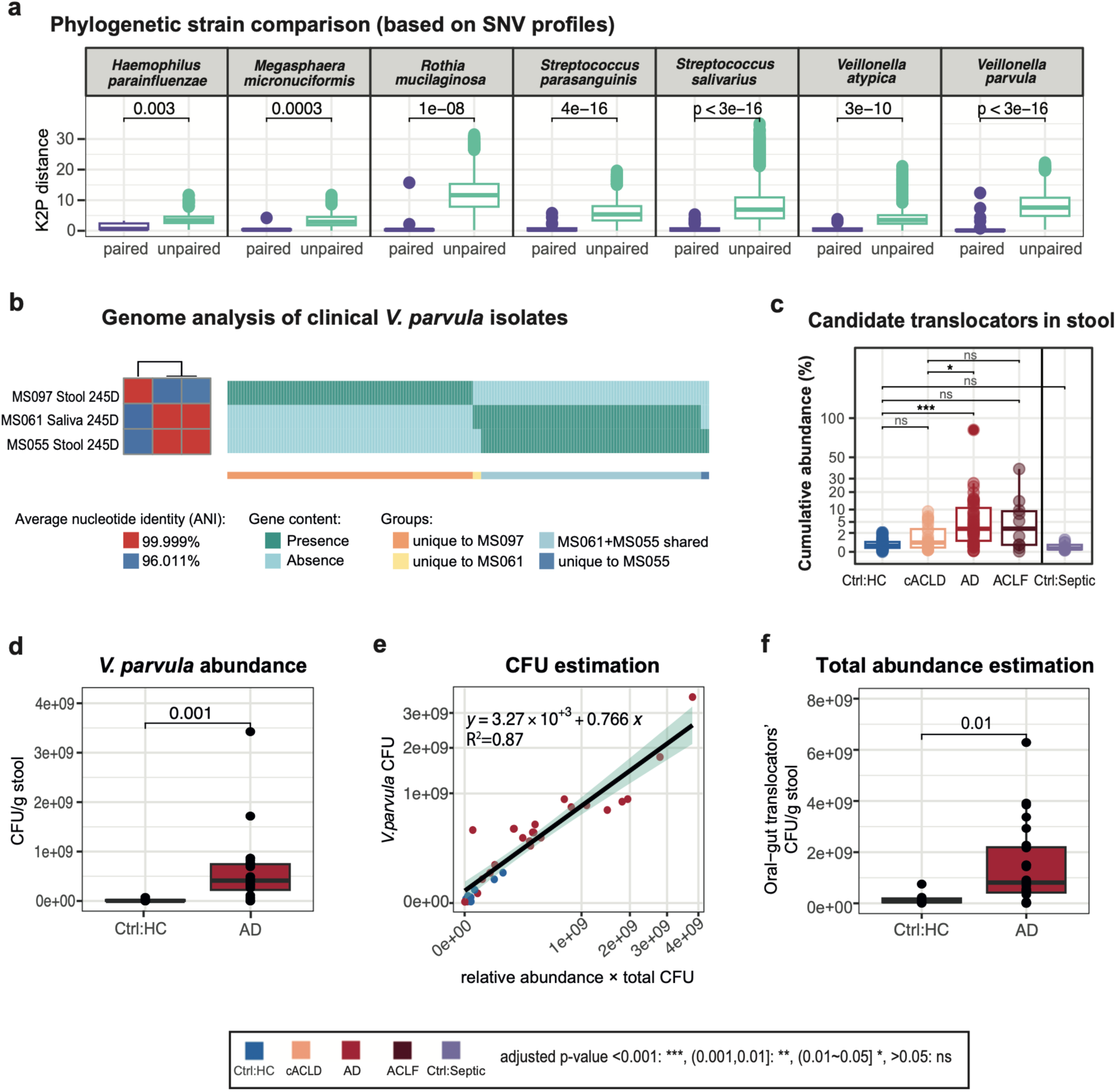
Identification of potential oral-gut translocators and their association with clinical ACLD parameters. **(a)** Comparison of phylogenetic distance between strains from paired versus unpaired faecal and saliva samples. The y-axis shows Kimura 2-Parameter (K2P) genetic distance estimates based on clade-specific marker genes (StrainPhlAn). **(b)** Genome analysis of clinical *V. parvula* isolates from paired faecal and saliva samples of an AD patient (patient-id 245D, AD: acutely decompensated ACLD). Genomes from three representative strains are shown, including phylogenetic relationship (left) based on genome-wide average nucleotide identity (ANI) and gene presence/absence patterns (right, PanPhlAn), where genes shared by all three isolates were omitted (1,539 genes shared by all three isolates, 12 unique to MS061, 12 unique to MS055, 353 unique to MS097, 316 uniquely shared by MS061 and MS055). **(c)** Cumulative relative faecal abundance of candidate translocators stratified by disease severity (Wilcoxon, ns = not significant, p-value adjusted with BH). **(d)** Absolute abundance of *V. parvula* per disease group based on qPCR analysis (Wilcoxon). **(e)** Comparison of quantitative *V. parvula* abundances (qPCR with nifJ primer, y-axis) and estimated *V. parvula* abundance based on total bacterial load multiplied by relative *V. parvula* abundance from MetaPhlAn (x-axis). A linear regression was fitted with R=0.87. **(f)** Absolute abundance of oral-gut translocators was estimated based on their cumulative relative abundance (MetaPhlAn) multiplied by total bacterial load (qPCR, universal 16S primer). A significant increase in the abundance of oral-gut translocators was observed in AD patients (Wilcoxon).

Next, viable strains were selectively isolated from patient samples based on the computationally identified oral-gut translocation signals. Next-to-identical isolates for *V. parvula, V. dispar, S. gordonii* and *S. parasanguinis* were obtained from paired saliva and faecal samples with average nucleotide identities (ANI) > 99.98%. We focused our subsequent followup investigation on the most frequent translocator in ACLD patients: *Veillonella parvula*. *V. parvula* paired oral and faecal isolates showed an ANI of 99.999%, shared 99.4% of genes (MS061 and MS055, **Figure 2b**) and exhibited highly similar growth patterns under various growth conditions (**Figure S1f**). Interestingly, an additional *V. parvula* strain was isolated from the same faecal sample with substantially higher strain heterogeneity compared to the next-to-identical isolate-pair (MS061 and MS055), showing a genome-wide ANI of 96% and 353 unique genes (**Figure 2b**). This strain heterogeneity motivated our subsequent investigation of common gene signatures among translocating strains.

### Increased oral-gut translocation in ACLD is associated with disease severity and intestinal barrier dysfunction

The cumulative abundance of these oral-gut translocators significantly increased in the gut of ACLD patients (**Figure 2c**), where in particular, *V. parvula*, *V. atypica*, *S. parasanguinis* and *Megasphaera micronuciformis* were significantly enriched in cirrhosis (**Figure S2b**). While some of these species were also commonly detected in healthy controls (e.g. *S. salivarius* was present in >40% of paired healthy samples) and sepsis patients, their faecal abundance was very low (average total abundance of 0.4% and 0.2%, respectively). Notably, faecal abundances of *V. parvula*, *V. atypica*, *V. dispar*, *V. infantium* and *M. micronuciformis* showed positive correlations with ACLD severity indices (**Figure S2a**), including Child-Pugh and MELD scores. Both Child-Pugh and MELD scores are important clinical measurements of cirrhosis severity used to evaluate need for orthotopic liver transplantation and determining disease prognosis [40]. These analyses suggest that increasing oral-gut translocation may reflect ACLD progression.

We next investigated whether these oral bacteria increased in absolute or only relative abundances in the faeces of AD patients. Comparable levels of total bacterial biomass were found in AD patients and healthy controls (**Figure S2c**), indicating a quantitative increase of oral bacteria in AD. We also directly confirmed quantitative increases of *V. parvula* in AD patients (**Figure 2d**). Importantly, *V. parvula* abundance estimation based on total microbial load and MetaPhlAn relative abundances agreed with the qPCR-based *V. parvula* measurements (**Figure 2e**). This confirms that species’ absolute abundances can be inferred based on MetaPhlAn relative abundance information in combination with total biomass estimation. Therefore, we also estimated overall absolute abundances for all oral-gut translocators (**Figure 2f**), confirming an absolute increase in the abundance of the identified oral-gut translocators in AD patients.

Next, we checked for associations of oral-gut translocation with intestinal barrier dysfunction using plasma levels of FABP2. FABP2 is a cytosolic fatty acid binding protein uniquely expressed in the small intestinal epithelium [41–43] and increased FABP2 levels in peripheral circulation indicate damage to the intestinal epithelium, which may in turn lead to increased epithelial permeability [44]. Plasma FABP2 levels were significantly increased in the ACLD groups, in particular AD patients (**Figure S3a**). Notably, all identified oral-gut translocators showed significant positive correlations of their faecal abundance with plasma FABP2 levels (**Figure 3a**). Intestinal barrier impairment has been hypothesised to drive cirrhosis-associated immune dysfunction, increase the risk of infection [45, 46] by allowing bacteria and their products to enter systemic circulation and propagate hepatic inflammation via direct portal venous inflow [47, 48]. Our results indicate a potential link between ectopic gut colonisation of oral bacteria with ACLD progression and intestinal barrier impairment.

**Figure 3:**
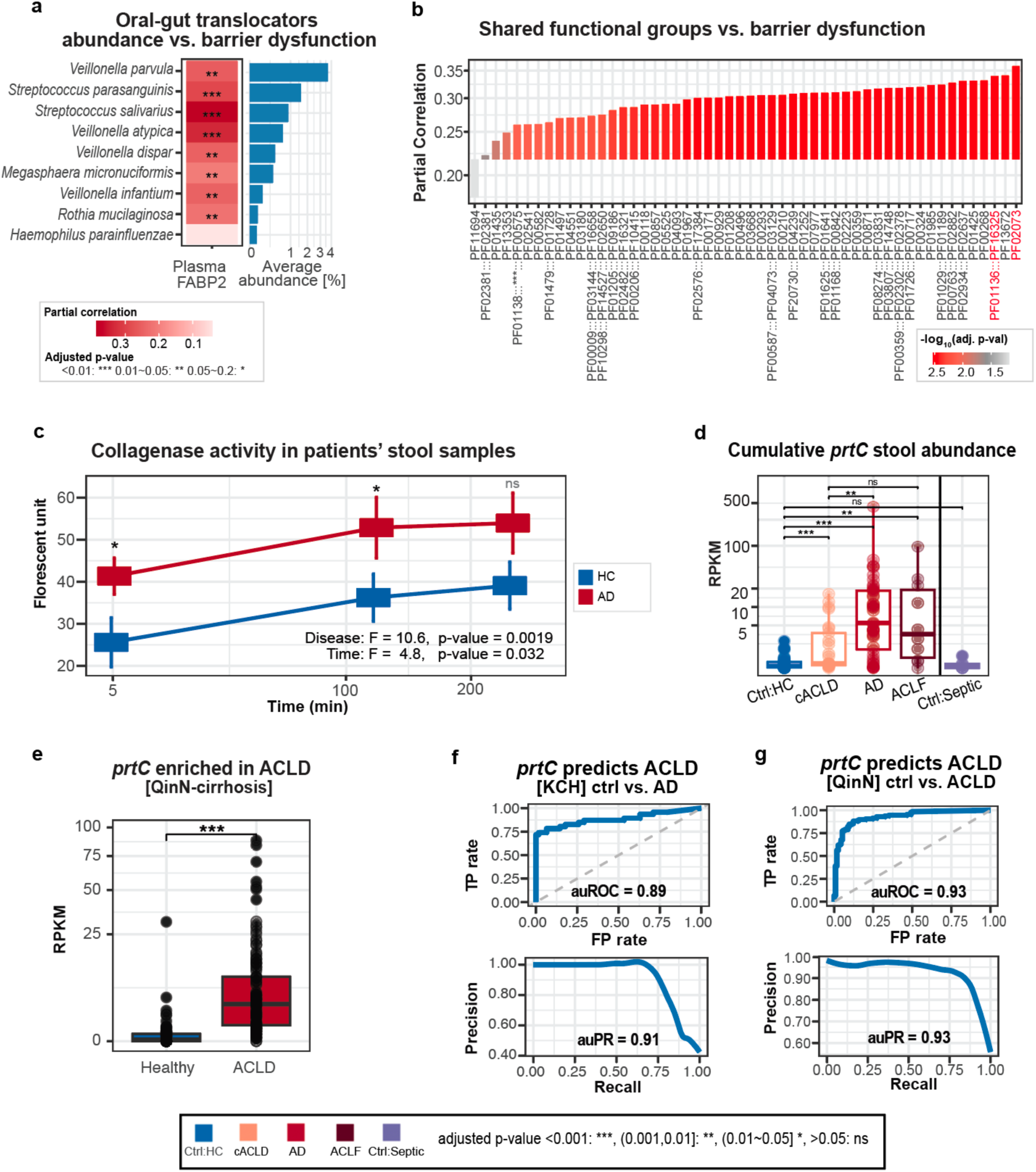
Functional analysis reveals putative microbial mechanisms resulting from oral-gut translocation involved in damage to the intestinal barrier. **(a)** Faecal abundance of oral-gut translocators is positively correlated with FABP2 levels (partial correlation, Spearman, corrected for age, gender antibiotic usage, p-values were adjusted using Benjamini Hochberg [BH], adjusted p-value <0.01:***, 0.01∼0.05:**, 0.05∼0.2:*). Species were ordered by average stool abundance in AD patients (shown on the right). **(b)** Stool abundances of functional groups unique to translocators were positively correlated with FABP2 levels (partial correlation, Spearman, corrected for age, gender antibiotic usage, p-values were adjusted with BH). Proteinases are highlighted in red (x-axis), including thermophilic metalloprotease (M29, PF02073) and collagenase-like proteinase (prtC, PF01136:::PF16325). PF01138:::***:::PF00575 = PF03725:::PF03726:::PF01138:::PF03725:::PF00013.” **(c)** Collagenase activity in faeces for AD patients (red) and healthy controls (blue). X-axis shows incubation time (min) and y-axis fluorescences. Error bars indicate standard error (ANOVA; disease: F=10.6, p-value=0.0019; time: F=4.8, p-value=0.032). **(d)** Cumulative *prtC* abundance in faeces compared across disease groups (Wilcoxon, p-value adjusted with BH). **(e)** Comparing total *prtC* faecal abundance (RPKM, y-axis) per disease group in a replication cohort (QinN-cirrhosis, N_healthy_=114, N_cirrhosis_=123, Wilcoxon). Species-level *prtC* homologs were identified using blastn on the corresponding gene catalogue (>95% identity, >90% coverage). At least one *prtC* gene was identified for each translocating species and their cumulative abundance was significantly increased in the ACLD group (Wilcoxon, p-value< 2.22e^−16^, mean abundance increased 20.6-fold compared to the healthy group). **(f)** ROC and PR curves for disease predictions (AD vs controls) based on prtC faecal abundance in our cohort (KCH, auROC=0.89, auPR=0.91) and **(g)** analogously for an independent replication cohort (ACLD vs controls, QinN, auROC=0.93, auPR=0.93).

### Intestinal barrier impairment in ACLD patients is associated with a bacterial collagenase gene shared among oral-gut translocators

To further investigate aberrant host-microbial interactions involving intestinal barrier impairment, we next looked for shared genes across oral-gut translocating species that are not typically present in the gut microbiota. For this, we grouped genes from the gene catalogue into functional groups (FGs, **Figure S3b**), where genes encoding the same functional domains in the same order were grouped together, and mapped these FGs to the MSPs of the oral-gut translocators. In total, 288 FGs were commonly shared by MSPs assigned to oral-gut translocating species (>80% prevalence). To filter out housekeeping genes and FGs normally present in the gut, we compared these 288 FGs against FGs from the core genomes of typical prevalent gut commensals. MSPs with mean abundance >5% in healthy faecal samples were chosen for this comparison, including *Prevotella copri*, *Bacteroides uniformis* and *Faecalibacterium prausnitzii*. After this filtering step, 52 FGs remained, which potentially encode mechanisms essential for oral-gut translocation. We next associated faecal abundances of these 52 FGs with plasma FABP2 levels (**Figure 3b**). Two of the top FGs were related to proteolytic activity, including a thermophilic metalloprotease (M29) and a peptidase with U32 protein. The latter belongs to a family of proteinases with enzymatic activity to degrade collagen. A well-studied case is protease C (*prtC*) in *Porphyromonas gingivalis*, a pathogenic bacteria associated with periodontal diseases, that degrades human type-I collagen [49], induces pro-inflammatory cytokines (IL-1a, IL-8, and TNFa) and leads to tissue loss in periodontal disease [50]. *PrtC* homologs shared by oral-gut translocators showed similar domain organisation compared to the *P. gingivalis* collagenase encoding gene (P33437), characterised by a peptidase U32 domain followed by an additional peptidase U32 C-terminal domain (**Figure S3c**). While sequence similarity was relatively low (40%), the predicted structures (AlphaFold2) were highly similar compared to the experimentally characterised collagenase of *P. gingivalis* (RMSD<4.7Å, **Figure S3d**), suggesting potential for its human type-1 collagen degradation ability.

To gain additional evidence for the role of collagen degradation in ACLD pathophysiology, we measured collagenase activity in patient faecal samples. Significantly higher collagenase activity was observed in AD patients compared to controls (p < 0.0019, F=10.6 **Figure 3c**). Type-I collagen is critical for structural tissue integrity [51] [52], therefore higher collagenase activity suggests increased collagen degradation, which potentially leads to increased barrier dysfunction. Overall, this suggests that ectopic gut colonization by oral bacteria introduces microbiome functional shifts (i.e. collagenase activity) that may contribute to disease-associated gut barrier impairment in ACLD.

### Faecal *prtC* abundance is a robust biomarker for ACLD

Total *prtC* abundance was significantly increased in both faecal and saliva samples in ACLD, particularly in AD patients (mean *prtC* abundance showed a 94.8-fold increase in faecal and 1.8-fold increase in saliva AD samples compared to healthy controls, **Figure 3d + S4a**). We verified this cirrhosis-associated *prtC* gene signal using several independent replication cohorts (**Figure S4b**). In the ACLD replication cohort (QinN-cirrhosis), we identified at least one *prtC* homolog at species level for all oral-gut translocators (>95% identity, >90% coverage). In contrast, in the IBD cohort (HMP2-IBD) only one *prtC* gene from *V. parvula* was present. Overall, *prtC* gene abundance was significantly increased in QinN-cirrhosis (**Figure 3e**, 20.6-fold increase of mean abundance in cirrhosis group) but not in HMP2-IBD (**Figure S4c**) or in healthy populations (500FG and LifeLinesDEEP, **Table 2**). Together, these results suggest that oral-gut translocation in ACLD leads to ectopic gut colonisation of collagen-degrading bacteria, which has the potential to damage intestinal connective epithelial tissue leading to increased gut barrier dysfunction.

**Table 2:** Bioinformatic analysis and followup experiments. **(a)** Closest homologs matching the *prtC* genes based on the UniProt database for structure prediction (blastx, AlphaFold). **(b)** Summary of cirrhosis prediction based on stool *prtC* gene abundance (detection limit: > 0.5 RPKM).

Using the cumulative abundance of the *prtC* gene from oral-gut translocators as a feature, we were able to reliably distinguish the control groups (ctrl:HC and ctrl:septic) from the ACLD groups (auROC=0.84, auPR=0.90, **Figure S4e**). The predictive power increased for AD (auROC=0.89, auPR=0.91, **Figure 3f**). These signals were confirmed in two ACLD replication cohorts (QinN-cirrhosis: auROC=0.93, auPR=0.93, **Figure 3g**; CLD-Sole2021, **Figure S4f**). In previous studies, ACLD prediction models based on the abundance of 19 species combined with patient age only achieved an auROC of 0.86 in the QinN-cirrhosis cohort [12]. We also verified the specificity of our model in two healthy cohorts (500FG [53] and LifeLinesDEEP [54, 55] with 471 and 1,135 participants, respectively), where the model achieved 100% specificity in 500FG and 92.5% specificity in LifeLinesDEEP. These analyses identify faecal *prtC* gene abundance as a strong and robust predictor of ACLD.

### Clinical isolates of oral-gut translocators exacerbate intestinal barrier dysfunction and hepatic fibrosis in a CCl_4_ mouse model

We next investigated the ability of bacterial oral-gut translocators to exacerbate gut barrier impairment *in vivo* using *prtC*-encoding *Veillonella parvula*, *Veillonella dispar* and S*treptococcus parasanguinis* strains, which were isolated from ACLD patients’ saliva (**Table S1**). Mice were treated twice a week with carbon tetrachloride (CCl_4_) to induce hepatic fibrosis. After the first two weeks, one group of mice continued with the same CCl_4_ treatment (CCl_4_ group) and the other group additionally received a cocktail of oral isolates for three consecutive days per-week (CCl_4_+isolates group). At the end of week six, mice were sacrificed for sampling and downstream analyses (**Figure 4a**). While CCl_4_-treated mice generally showed a significant increase in colonic albumin compared to non-treated controls (p=0.036 and p=0.005, respectively), the highest albumin levels were observed for the CCl_4_+isolates group (**Figure S4g**) indicating increased gut barrier disruption [56, 57]. Immunofluorescent images provided additional evidence for disrupted epithelial junction architecture: tight junction proteins (E-cadherin and occludin) in the CCl_4_+isolates group were relocated to the cytoplasm (**Figure 4b+c**), indicating a loss of membrane localisation [58]. These signals highlight a role for oral-gut translocating strains in disrupting gut barrier integrity.

**Figure 4:**
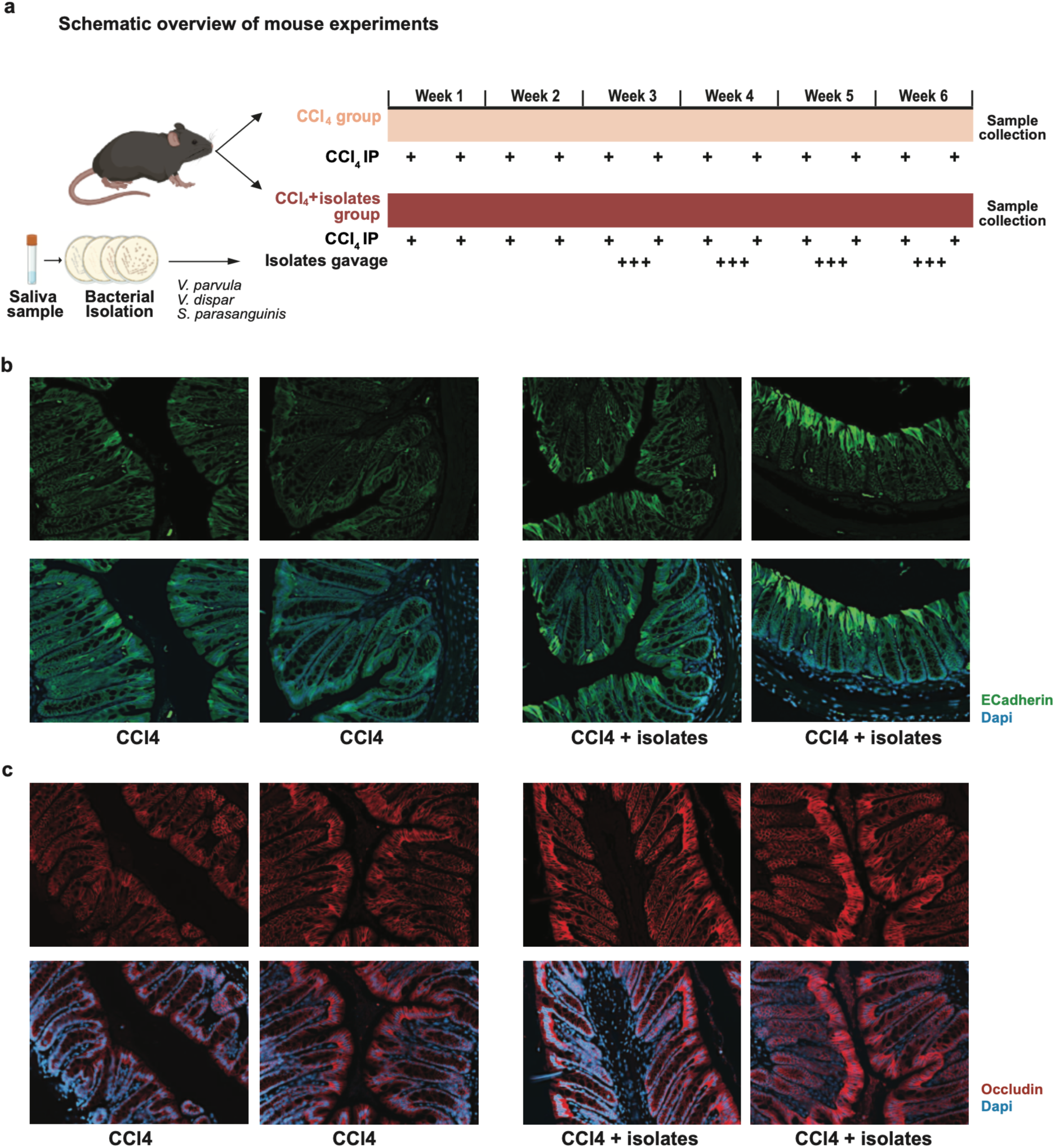
Clinical isolates linked to oral-gut translocation induce intestinal barrier dysfunction in a mouse model of fibrosis. **(a)** Experimental overview: Strains associated with oral-gut translocation were isolated from patients’ saliva, including *Veillonella parvula*, *Veillonella dispar* and *Streptococcus parasanguinis* (**Table S1**). C57BL/6 mice were treated i.p. with CCl_4_ twice a week for six weeks. After two weeks, one group of mice received oral gavage of patient saliva isolates 3x per week for 4 weeks (CCl_4_+isolates). Mice were sacrificed 2 days after the last CCl_4_ administration and samples were collected for analysis. **(b)** E-cadherin staining (green) highlighting adherens junctions along the epithelial lining. **(c)** Occludin staining (red) marking tight junctions between epithelial cells. Nuclei are counterstained with DAPI (blue) to provide structural context. Images are shown for both treatment groups (CCl_4_ and CCl_4_+isolates) to illustrate differences in junctional protein localization.

In addition, we also observed exacerbated hepatic and intestinal fibrosis and small intestinal bacterial overgrowth. Collagen deposition in the liver was observed for both the CCl_4_ and CCl_4_+isolates group (**Figure S5a**). Importantly, the fraction of liver fibrotic tissue was significantly increased in the CCl_4_+isolates group (**Figure 5a**), indicating an exacerbation of hepatic fibrosis. Furthermore, the degree of liver fibrosis was correlated with colonic albumin levels (tau=0.35, p-value=0.08, Kendall’s rank correlation, **Figure 5b**), suggesting a potential link between gut barrier dysfunction and liver fibrosis exacerbation. Intestinal barrier impairment is recognized as a driver of hepatic fibrogenesis in ACLD and preserving gut integrity can attenuate liver fibrosis [59]. Intestinal fibrosis was also quantified for the mucosa, muscularis mucosae, submucosa and muscularis regions (details in **Figure S5b** and method section). Fibrosis-related disruption of mucosal architecture and barrier integrity is increasingly recognized as a key contributor to the development and progression of chronic liver disease [60]. Indeed, the CCl_4_+isolates group also showed a significant increase in intestinal fibrosis (**Figure S5c**), suggesting a potential role of these clinical isolates in exacerbating both intestinal and liver fibrosis. Additionally, total bacterial load in the distal ileum (P3, **Figure S5d**) was increased, revealing ileal bacterial overgrowth in the CCl_4_+isolates group (**Figure 5c**). Together, these results provide *in vivo* evidence for a causal role of oral bacteria in ACLD progression, including exacerbated gut barrier impairment and hepatic and intestinal fibrosis.

**Figure 5:**
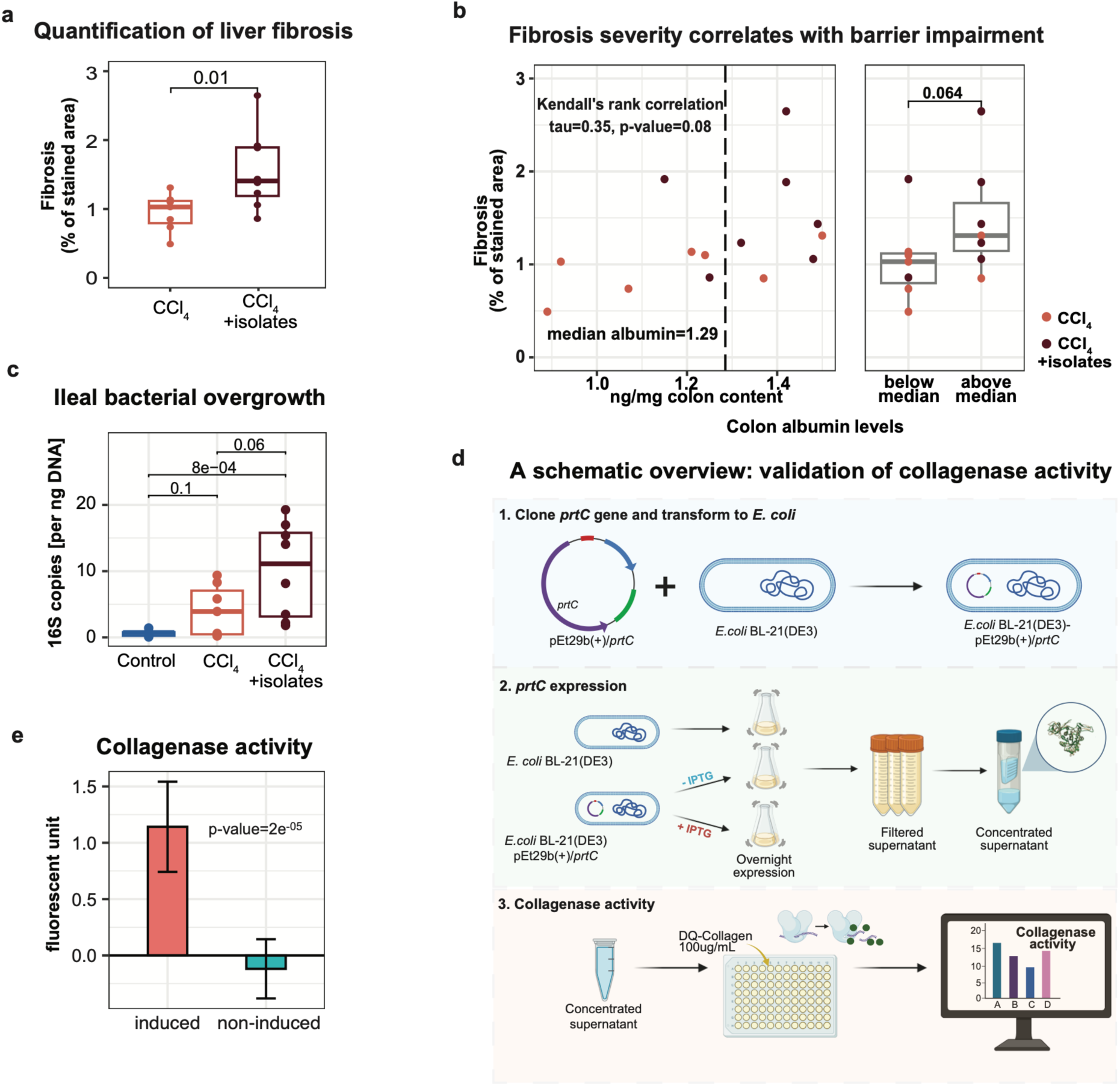
Oral clinical isolates with collagenase activity exacerbate liver fibrosis in mice. **(a)** Quantification of liver fibrosis was calculated as the percentage of area stained by Sirius red. The y-axis refers to the estimated fraction of fibrous tissue for CCl_4_ or CCl_4_+isolates, respectively (one-sided Wilcoxon). **(b)** Fibrosis severity was positively correlated with intestinal barrier impairment (tau=0.35, p-value=0.08, Kendall’s rank correlation). Left: X-axis indicates albumin level (ng/mg colonic content) and y-axis represents the percentage of fibrosis in liver tissues. The dashed line shows median albumin level across all samples and color indicates different mouse groups. Right: Samples were stratified into two groups based on albumin levels below or above the median. Fibrosis severity was compared between both groups (p-value=0.064, one-sided Wilcoxon). **(c)** Comparison of ileal bacterial overgrowth among control, CCl₄-treated, and CCl₄+isolates-treated mice. The y-axis indicates bacterial load, quantified as 16S rRNA gene copies per nanogram of DNA. **(d)** Schematic overview of experimental validation of collagenase activity of the *V. parvula prtC* gene, including transformation of *E.coli* BL21(DE3) with pET29b(+)/*prtC, prtC* expression and supernatant concentration and collagenase activity measurements. **(e)** Comparison of collagenase activity of induced (+IPTG) and non-induced (-IPTG) *E.coli* BL21(DE3)-pET29b(+)/*prtC* (Wilcoxon test).

### Experimental validation of *V. parvula prtC* collagenase activity

Lastly, we confirmed collagenase activity of the bacterial *prtC* gene identified in the oral-gut translocators. We focused on the *prtC*-encoding collagenase from *Veillonella parvula*, the most frequent translocator. The gene was cloned into *E. coli* BL21 (DE3) using the pEt29b(+) expression system (**Figure 5d**). Expression was induced and resulted in a distinct band corresponding to the expected molecular weight of the recombinant protein, which was absent in the supernatant of the non-induced strain (48kDa, **Figure S5e**). Importantly, functional assay confirmed an active and functional collagenase protein (**Figure 5e**).

Taken together, these results support the role of oral-gut translocation of bacterial strains, such as *prtC*-encoding *Veillonella parvula*, and gives mechanistic insights into how this may exacerbate disease progression, including gut barrier disruption through collagenase activity in ACLD patients. This in turn may promote further microbial translocation along the gut-liver axis (leaky gut hypothesis) and the exacerbation of liver and intestinal fibrosis. Importantly, bacterial products in peripheral circulation, such as endotoxins [61–63], have been associated with immunological alterations and inflammation in ACLD patients [61]. Overall, restoration of the intestinal barrier and modulation of the oral microbiome may be a promising avenue for preventing ACLD progression and the development of subsequent clinical complications, including hepatic decompensation and extra-hepatic organ failure.

## Discussion

In this study, we investigated the role of oral-gut translocation in ACLD pathobiology. While increases in bacteria typical for the oral cavities in faecal samples of ACLD patients were observed previously [12, 23], it was unknown whether these bacteria originate from the oral cavities (translocation), if they ectopically colonize the gut (engraftment) and how they are involved in ACLD progression (disease contribution). Our results show that next-to-identical bacterial strains are present in paired saliva and faecal samples of ACLD patients and that the identified oral-gut translocators increase in absolute abundances, providing evidence for gut engraftment. Importantly, we confirmed their disease involvement *in vivo*, where oral gavage with patient isolates exacerbated intestinal barrier impairment and gut and liver fibrosis in mice, providing mechanistic insights into how the oral microbiome may contribute to ACLD pathobiology.

Using microbiome strain and functional profiles, we bioinformatically identified putative microbial disease mechanisms, including a collagenase-like proteinase (*prtC*), which we linked to collagen degradation. *PrtC* homologs were uniquely encoded by oral-gut translocators but absent in commensal gut microbes. While structurally similar proteins are linked to epithelium damage in periodontal disease [64], translocation-related consequences for the gut were unknown. Complementary evidence supports the relevance of this bacterial gene in ACLD, including (1) faecal *prtC* abundance, which showed a strong positive correlation with markers of intestinal barrier damage (FABP2), (2) experimental validation of bacterial *prtC* collagenase activity using a recombinantly expressed *V. parvula prtC* gene and (3) higher collagenase activity in faecal samples of ACLD patients. Previously, elevated collagenase activity was also reported for mouse faecal supernatant in the context of hepatocellular carcinoma (HCC) linking *Klebsiella pneumonia* to carcinogenesis [65]. Our study provides clear evidence for a role of oral-gut translocation in collagenase increase and further motivates the investigation of the role of the oral microbiome in HCC. Importantly, we were also able to show that the faecal abundance of a single bacterial gene (i.e. *prtC*) can serve as a robust biomarker to distinguish between ACLD and healthy individuals (auROC=0.89, auPR=0.91, sensitivity=0.67, specificity=0.89), putting it on par with current established non-invasive diagnostic methods used in the clinic, such as transient elastography (sensitivity=0.61, specificity=0.95) [66] and magnetic resonance elastography (sensitivity=0.86, specificity=0.85) [67].

Intestinal barrier impairment was further investigated *in vivo* in mice gavaged with oral patient isolates, which exhibited increased colon albumin levels, small intestinal bacterial overgrowth (SIBO) and endocytosis of tight junction proteins (E-cadherin and occludin), which moved from the basal compartment to the cytosol of gut epithelial cells. These *in vivo* findings are consistent with our observations in patients, where oral bacteria increased in absolute abundance in the faeces of ACLD patients, which was in turn associated with impaired gut barrier function (increased plasma FABP2 levels). Interestingly, mice gavaged with patient isolates showed other characteristics that were previously observed in patients with chronic intestinal inflammation, including gut fibrosis, which involved excessive extracellular matrix deposition and scarring that impaired intestinal function and barrier integrity by altering motility and mucosal architecture [68] [69]. These gut changes facilitate SIBO and endotoxin translocation. Once they enter the portal vein, endotoxins activate Kupffer and hepatic stellate cells, driving hepatic inflammation and fibrogenesis [70]. Furthermore, the leaky gut hypothesis suggests that increased intestinal permeability [44] leads to increases of bacterial products in systemic circulation, which may be involved in hepatic inflammation and subsequent disease progression [71, 72]. Our study provides new evidence that links oral-origin microbes to a leaky gut in ACLD, implicating specific collagenase degrading bacterial genes and strains from the oral cavity in intestinal barrier impairment and hepatic fibrosis progression.

We uncovered several new important insights in our study, however, there are also limitations that apply and will motivate subsequent follow-up work. Our findings are based on a cross-sectional, single-centre cohort with patients located in the United Kingdom. We used 5 replication cohorts (USA, Europe and China) to ensure that our results are applicable to a wide spectrum of ethnicities and populations. Future work will include longitudinal sampling to measure sequential changes in oral-gut translocation and in response to specific therapies. Further, parenteral antibiotic treatment is commonly required for hospitalised ACLD patients and oral antibiotics are often used as prophylaxis in cACLD patients. Samples were collected as close to admission as possible (<48h after commencement of antibiotic therapy) to limit antibiotic effects on the microbiome. Our healthy controls differed from the ACLD groups in age, antibiotic usage, and PPI exposure, as it is often the case in patient cohort studies. To mitigate this, we included a second positive disease control group, patients with sepsis but no underlying ACLD, which share similar clinical characteristics (ACLD vs. Ctrl:septic: age 54.3±11.5 vs. 56.7±13.4, antibiotic exposure 62.8% and 92.9% and PPI usage of 53% and 36%, **Table 1**). Importantly, no increase in the abundance of the identified oral-gut translocators was observed in the sepsis group. Although these sepsis patients also showed increased similarity of saliva and faecal samples compared to healthy controls, this was largely due to *E. coli* and *Enterococcus faecium* overgrowth. Because both species are commonly detected in the gut and absent from healthy oral cavities, they were not considered oral–gut translocators. Furthermore, while we implicated a combination of oral-gut translocating strains in liver disease exacerbation in mice, future work will be required to evaluate the effect of individual strains and genes, which necessitates the development of genetic tools for these species and the addition of germ-free mouse models. Lastly, other potential confounding factors, such as oral hygiene, dental health and nutrition, may contribute to changes in the microbiome but were not recorded as part of this study.

In summary, we combined data-driven computational predictions from multi-omics data with *in vitro* and *in vivo* experimental validation of host-microbial interactions to provide novel insights into the origin and consequences of oral-gut translocation in ACLD. Linking oral-gut translocation and in particular the oral microbiome to ACLD pathobiology has the potential to facilitate the development of new therapeutic approaches. Re-establishing the commensal gut microbiome may also be a promising avenue to counteract ectopic gut colonization as already explored for *Enterobacteriaceae* infection [73]. Importantly, similar mechanisms may be at play in other chronic inflammatory diseases, as an enrichment of microbes typical for the oral cavities in the gut was also identified in IBD, colorectal cancer and rheumatoid arthritis [31]. Our study design and computational strategy provides a powerful framework to investigate and identify microbiome-derived mechanisms in these disease states, in particular in the context of bacterial translocation involving the gastrointestinal tract.

## Acknowledgements

The authors are grateful to all patient participants and healthy volunteers for agreeing to take part in the ‘Gut-Liver Axis in Liver Disease & Transplantation’ study, which was adopted to the National Institute for Health Research (NIHR) Clinical Research Network (CRN) portfolio, supporting participant screening and recruitment by the Liver Research and Anaesthetics, Critical Care, Emergency and Trauma Teams at King’s College Hospital NHS Foundation Trust. This work was further supported by a grant from the Deutsche Forschungsgemeinschaft (DFG, German Research Foundation) [426120468 to MeS] and TUM Innovation Network NextGenDrugs (MeS) funded under the Excellence Strategy of the Federal Government and the Länder. Further laboratory assays and personnel were also funded by the Foundation for Liver Research (Registered charity number: 268211/1134579) and a generous donation from a Liver Service User to the King’s College Hospital Institute of Liver Studies and Transplantation Charitable Research Fund, King’s College Hospital Charity (Registered charity number: 1165593). VCP was supported by the NIHR South London CRN Strategic Greenshoots Funding scheme. The work was also supported by the Science for Life Laboratory (SciLifeLab) and KTH-Royal Institute of Technology and by the Engineering and Physical Sciences Research Council (EPSRC), EP/S001301/1 and Centre for Host-Microbiome Interactions at King’s College London. NB and MMG were supported by the BBSRC Institute Strategic Programme Food Microbiome and Health BB/X011054/1 and its constituent project BBS/E/F/000PR13632 (NB) and the BBSRC Core Capability Grant BB/CCG1860/1 at the Quadram Institute Bioscience. The authors acknowledge support from the National Genomics Infrastructure in Stockholm funded by Science for Life Laboratory, the Knut and Alice Wallenberg Foundation and the Swedish Research Council, and SNIC/Uppsala Multidisciplinary Center for Advanced Computational Science for assistance with massively parallel sequencing and access to the UPPMAX computational infrastructure, under project number Project SNIC 2020-5-222, SNIC 2019/3-226, SNIC 2020/6-153, SNIC 2021/6-89, SNIC 2021/5-248 and SNIC 2021/6-242. SL was supported by GIST Research Institute (GRI) GIST-MIT research Collaboration grant by the GIST in 2022, and Basic Science Program (NRF-2021R1C1C1006336) & the Bio & Medical Technology Development Program (2021M3A9G8022959) of the Ministry of Science, ICT through the National Research Foundation, Korea. We also thank the LifeLines-DEEP participants and the Groningen LifeLines staff for providing and sharing their valuable data.

## Author contributions

SJ analysed and interpreted the data and wrote the manuscript. AC performed the gene structure analysis, collagenase assay on patient samples and recombinant *V.p. prtC* expression, and wrote the manuscript. DW isolated and characterised clinical strains from patient samples. BA undertook sample processing, saliva and faecal DNA extractions and raw data handling, and measured plasma FABP2, as did MM. MaS and MM undertook data curation. AZ recruited patients, assisted with biological sampling and undertook data collection. SL assisted with raw microbiome data curation. DW and ML performed absolute abundance quantifications of microbial biomass and *V. parvula*. ER performed bioinformatic analysis and pipeline development for metagenomic data processing. SS and AM secured funding for initial metagenomic sequencing and were involved in initial project management. NB planned and supervised the mice experiment and performed IF imaging. MMG performed the mouse experiments and analyses, including tissue staining, quantification of fibrosis, 16S qPCR for SIBO, *in vivo* experiments and ELISAs (albumin). VCP conceptualised, designed, supervised and project managed the clinical study and served as the clinical study Principal Investigator, coordinated and secured funding and resources for study participant recruitment, data acquisition and experimental assays (DNA extractions, metagenomic sequencing and FABP2), and edited the original manuscript with MeS. MeS designed, supervised and coordinated the bioinformatic analysis and *in vitro* follow-up experiments, obtained funding for this part of the work and additional metagenomic sequencing, and wrote the manuscript. All authors provided editorial input and approved the final version of the manuscript.

## Methods

### Data and code availability

Metagenomic reads will be made available upon acceptance of the paper. Processed metagenomic profiles, including MetaPhlAn3 taxonomic profiles, MSP abundances and MSP taxonomy annotation can be found in **Table S2**. All additional data related to this work is also included in **Table S2**, including clinical metadata, FABP2 measurements and disease scores. The pipeline used for metagenomic quality control, reference-based microbial profiling and gene-centric analyses is available at https://github.com/schirmer-lab/metagear-pipeline.

### Study participants & biological sampling

Patients were consecutively recruited at King’s College Hospital after admission to the ward or from the hepatology out-patient clinic. The study was granted ethics approval by the national research ethics committee (12/LO/1417) and local research and development department (KCH12-126) and performed conforming to the Declaration of Helsinki. Patient participants, or their family nominee as consultees in the case of lack of capacity, provided written informed consent within 48 hours of presentation. Patients were managed according to standard evidence-based protocols and guidelines [75]. Patient and Public Involvement and Engagement was undertaken with a patient advisory group who partnered with us to determine acceptability of the study, provided their perspective on study design, informational material, measures to minimise participation burden and agreeing on a dissemination plan of the findings.

Patient participants were stratified into and phenotyped according to clinically relevant groups based on the severity and time course of their underlying cirrhosis, degree of stability and hepatic decompensation, and presence and extent of hepatic and extrahepatic organ failure at the time of sampling. These groups were compensated ACLD (cACLD), acutely decompensated ACLD (AD) and acute-on-chronic liver failure (ACLF), with separately recruited healthy controls (Ctrl:HC) and septic patients with no underlying ACLD as an additional control group (Ctrl:Septic). AD was defined by the acute development of one or more major complications of cirrhosis, including ascites, hepatic encephalopathy, variceal haemorrhage, and bacterial infection. ACLF was defined and graded according to the number of organ failures in concordance with criteria reported in the CANONIC study [76] [77]. Main exclusion criteria included pregnancy, hepatic or non-hepatic malignancy, pre-existing immunosuppressive states, replicating HBV/HCV/HIV infection, and known inflammatory bowel disease (IBD).

Demographic, clinical, and biochemical metadata were collected at the time of biological sampling. Standard clinical composite scores used for risk stratification and prognostication included the Child-Pugh score [78] and model for end-stage liver disease (MELD) [79]. For patients with sepsis without chronic liver disease (Ctrl:Septic), the diagnosis of sepsis was based on the Sepsis-3 criteria [80] in which life-threatening organ dysfunction caused by a dysregulated host response to infection was evident, with organ dysfunction defined by an increase in the sequential (sepsis-related) organ failure assessment (SOFA) score of 2 points or more. The absence of chronic liver disease in this patient group was determined by a combined assessment of clinical history with biochemical and radiological parameters.

Healthy controls aged >18 years (n = 52) were recruited to establish reference values for the various assays performed. Exclusion criteria for healthy controls were body mass index <18 or >27; pregnancy or active breastfeeding, a personal history of thrombotic or liver disease; chronic medical conditions requiring regular primary or secondary care review and/or prescribed pharmacotherapies; or current use of anticoagulants, platelet function inhibitors, or oral contraceptives.

### Plasma FABP2 quantification

Plasma samples for FABP2 profiling and quantification were obtained within 24 hours of admission to hospital. Intestinal fatty-acid-binding protein-2 (FABP2) [42] [44] was quantified to serve as a gut-specific marker of intestinal barrier integrity, to assess whether these differentiated at the different stages of cirrhosis and in the Ctrl:HC cohort to define whether physiological or basal levels were detectable. FABP2 was quantified using the human FABP2/I-FABP Quantikine ELISA Kit (R&D Systems). All assays were conducted according to the manufacturers’ instructions. Optical densities were measured with a FLUOstar® Omega Absorbance Microplate Reader.

### Faecal and saliva sample acquisition

Faecal samples were obtained within 48 hours of admission to hospital and collected into non-treated sterile universal tubes (Alpha LaboratoriesTM) without any additives. Faecal samples were kept at 4°C without any preservative and were homogenised within 2 hours, pre-weighed into 200mg aliquots in Fastprep tubes (MP BiomedicalsTM). Saliva samples were obtained within 48 hours of admission to hospital and collected into non-treated sterile universal tubes (Alpha LaboratoriesTM) without any additives. A controlled passive ‘drool’ was performed by the study participant into a universal container repeatedly until at least 6mL of saliva was obtained. For patients that were intubated for mechanical ventilation, oro-pharyngeal suctioning of accumulating oral secretions were obtained. Saliva samples were kept at 4°C without any preservative and within 2 hours were homogenised, and measured into 1mL aliquots using sterile wide bore pipettes in Fastprep tubes (MP BiomedicalsTM), which were then centrifuged at 17,000 g for 10 minutes. The saliva supernatant was removed and stored separately from the remaining pellet. Faecal and saliva samples were stored at −80°C for subsequent DNA extraction and metabolite measurements.

### Metagenomic data generation for saliva and faecal samples

Metagenomic data ENA Project ID: PRJEB52891 was generated as part of Lee et al. [74]. Briefly, microbial DNA was extracted from stored faecal and saliva pelleted samples utilising a two-day protocol adapted from the International Human Microbiome Standards (IHMS) [81] [82]. A 200mg pre-weighed and homogenised aliquot was used for faeces, whilst for saliva, a post-centrifugation pellet was used. Processing of additional healthy control samples (Project ID: PRJNA1307628 *[will be made available upon acceptance of the paper])*: sample collection and storage was identical to the initial cohort samples (details in [74]). DNA for the additional faecal samples was extracted with the AllPrep® PowerFecal® DNA/RNA Kit Qiagen (Kit Catalog No. 80244) and for saliva samples the extraction kit (DNeasy PowerSoil Pro Kit (Qiagen, 47014)) was used according to the manufacturer’s instructions. DNA was subsequently sequenced (Illumina NovoSeq, paired-end mode, read length 2×150bp) with a targeted sequencing depth of 25 Gbp for both saliva and faecal samples.

### Quantification of absolute bacterial abundances using qPCR

In order to quantify total bacterial biomass and *V. parvula* absolute abundances, we performed qPCR assays following the protocol by Suzuki et al. [83] and Rojas-Tapias et al. [35]. Briefly, we first conducted serial dilutions of *Escherichia coli* (strain BL21 (DE3)) culture grown in mGAM medium and enumerated colony forming units (CFU) for each of the dilutions. Next, DNA isolation was performed using 6ml of *E. coli* culture applying Phenol-Chloroform-Isoamylalcohol as described by Harju et al. [84]. A universal 16S rRNA primer [83] targeting the V9 region was used and 2 ul of DNA were added to the mix of 2 ul of nuclease free water, 0.5 ul of forward and reverse primers (10 uM) and 5 ul of 2 x SensiFast SYBR mix (BioCat, Heidelberg, Germany). The assay was run according to the following setup: initial denaturation for 2 min at 95°C, 40 cycles of 3 step cycling at 95°C for 15 sec, 53°C for 15 sec and 72°C for 15 sec followed by melting at 95°C for 30 sec, 53°C for 30 sec, 60°C for 1 sec with final cooling at 40°C for 30 sec. The same procedure was repeated for the *V. parvula* patient isolate with *nifJ* gene specific primers introduced by Rojas-Tapias et al. [35]. Based on the computed Cp values, calibration curves were established as described by Yao et al. [85] to set ratios between Ct values and calculated CFUs further enabling estimations of CFU per-gram faeces (**Figure S2d+e**). Next, patient samples for which DNA was available were processed using the same qPCR setup and master mix with both V9 region 16S and *V. parvula* specific *nifJ* gene primers and compared to the calibration curve.

### Metagenomic data analysis

The metaGEAR pipeline was used for metagenomic quality control, reference-based microbial profiling and gene-centric analyses (https://github.com/schirmer-lab/metagear-pipeline). Briefly, raw metagenomic reads went through the following quality control pipeline to remove (1) sequencing adaptors, (2) low-quality reads and (3) human host contaminations. First, we applied trim_galore [86] for adaptor removal using the parameters “--paired --phred33 –quality 0 --stringency 5 --length 10”. Then, we applied KneadData (https://github.com/biobakery/kneaddata) to remove low quality reads and host contaminations. Low quality reads and adaptors were removed using the parameters “-- trimmomatic-options ‘SLIDINGWINDOW:4:15 MINLEN:50”. Human reads were removed by mapping against the human reference genome (hg38). In addition, tandem repeats were removed using the default “trf” option. The “--reorder” flag was applied to match qc’ed forward and reverse reads.

For reference-based microbial profiling and strain comparison, we first applied MetaPhlAn3 [37] to profile the microbial community based on clade-specific marker genes. Strain-level profiles were then generated for the species of interest based on (1) single nucleotide polymorphisms in the marker genes using StrainPhlAn3 [37] and (2) presence/absence profiling of gene families using PanPhlAn3 [37].

Assembly-based analyses for generating metagenomic assembled pangenomes included *de novo* assembly using MEGAHIT with default parameters [87] for each sample. Then protein coding genes were predicted using Prodigal with the “-p meta” flag [88] [89]. Incomplete genes were subsequently filtered with the “partial==00” flag. Afterwards, genes from different samples were merged and clustered to generate a gene catalogue using cd-hit-est [90] with parameters “-aS 0.9 -aL 0.9 -c 0.95” to group genes with sequence identity >95% and coverage >90%. Protein sequences were extracted for the representative sequences of each gene family and further grouped into a protein catalogue using cd-hit [91] [90] with parameters “-c 0.9 -aL 0.8 -aS 0.8” to group proteins with sequence identity >90% and coverage >80%. Abundance profiles for the clustered genes were generated using CoverM [92] contig (methods count and rpkm) using parameters "--min-read-percent-identity 95 --min-read-aligned-percent 75 --min-covered-fraction 20"

The protein catalogue was annotated with interproscan v5.47-82.0 [93] [94] using the parameter “-appl Pfam” to get domain level annotation. Functional group annotation was extracted for each representative sequence of the protein families by combining Pfam domain annotations according to their order for each protein. Afterwards, reads from each sample were mapped against the gene catalogue to obtain the respective abundance profiles. The Metagenomic Species Pangenomes (MSPs) were generated with MSPminer [95] with its default parameters. Taxonomy annotation for each MSP was predicted based on gtdb-tk (v2.3.0) [96] on its pangenomes and the abundance of each MSP was profiled by the median abundance of its core genes in each sample.

### Integration of reference- and assembly-based results

Many MSPs lacked species-level taxonomy annotations based on the gtdb-tk annotations. Therefore, we introduced a new taxonomy annotation approach for MSPs which additionally integrates information from the reference-based profiles. This approach is based on the assumption that each MSP will have a similar abundance profile as its corresponding referenced-based counterpart. Therefore, we inferred MSPs taxonomic information based on abundance correlations with reference-based abundance profiles (MetaPhlAn3). For each MSP, the species from the MetaPhlAn profile with the best abundance correlation was assigned as its taxonomic annotation.

### Definition of similarity score

The similarity score S is used to quantify the similarity between two samples at the microbial community level:

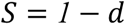

where d is the Bray-Curtis dissimilarity with values between 0 and 1. Therefore, S is also in the range between 0 and 1.

### Definition of dysbiosis score

We adapted the dysbiosis score introduced by Lloyd-Price *et al.* [34] to quantify the deviation of a given patient sample from the healthy control group. For our study, we calculated the dysbiosis separately for faecal and saliva samples. Specifically, for each body site, the dysbiosis score D for sample x is defined as the median Bray-Curtis distance to the samples from the healthy control group:

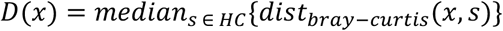

### Identification of candidate oral-gut translocators

First, we identified co-occurring species within each sample pair. For this, species were required to be present with a relative abundance of >0.1% in both sample types. Species were defined as candidate oral-gut translocators if they (1) frequently co-occurred in paired faecal and saliva samples (detected in at least 5 paired samples) and (2) were common members of the healthy oral microbiome (>1% abundance and detected in >20% of saliva samples). Overall, the number of co-occurring species was 26 (details in **Table S1**). Among these, 9 species were commonly detected in the oral samples, forming the final list of our candidate oral-gut translocators for the subsequent analyses. For each candidate translocator, we then compared the strain-level similarity between paired and unpaired samples using two different strategies: Phylogenetic similarity was quantified with the Kimura 2-Parameter (K2P) genetic distance based on marker gene alignment generated with StrainPhlAn3 [37]. The function distmat (https://www.bioinformatics.nl/cgi-bin/emboss/help/distmat) was used, which calculates the evolutionary distance between every pair of sequences in a multiple sequence alignment. Distances are expressed in terms of the number of substitutions per 100 bases. In addition, gene content similarity was compared, which was measured by the binary distance between the gene content of strains inferred with PanPhlAn3 [37].

### Associations of bacterial species with disease severity and plasma barrier dysfunction

The faecal abundance of oral–gut translocators was analyzed for associations with clinical indices, including disease severity (Child–Pugh and MELD scores) and barrier dysfunction (plasma FABP2). We used the pcor.test function in R to calculate partial correlations and corresponding p-values between each species and each clinical index. Associations were computed using a rank-based Spearman method, with age, gender, and antibiotic usage included as covariates. The detection limit for MetaPhlAn was set at 0.01%, with abundances below this threshold set to zero. Benjamini–Hochberg adjusted p-values were calculated separately for each clinical index, and results were visualized as a heatmap in which color indicates the partial correlation and asterisks denote significance levels (<0.01: ***, 0.01∼0.05: **, 0.05∼0.2: *).

### Multiple-testing correction for statistic analyses

#### Statistical analyses involving the five disease subgroups

For these analyses the healthy control group, ctrl:HC was compared to each of the disease subgroups, including cACLD, AD, ACLF and ctrl:Septic. In addition, the cACLD group was compared to the AD and ACLF to evaluate differences involved in disease progression. This results in six statistical tests in total. All p-values were correct for multiple testing (Benjamini–Hochberg) and adjusted p-values were reported in **Figure 1b–d**, **Figure 2c** and **Figure 3d**. For a subset of the supplementary figures the primary focus was on identifying changes between healthy controls (ctrl:HC) and the disease subgroups; here only four statistical comparisons were performed and corrected for multiple testing, including **Figure S1a**, **Figure S2b**, **Figure S3a** and **Figure S4a**.

#### Associations between oral–gut translocators and clinical indices

To test for associations between the nine species identified as potential oral-gut translocators with Child–Pugh, MELD scores and plasma FABP2 levels, respectively, 9 statistical tests were performed for each index. All p-values were corrected for multiple testing (Benjamin-Hochberg) and adjusted p-values are reported in **Figure S2a** and **Figure 3a**.

### Strain isolation

Saliva and stool strains were isolated on SK agar plates (as used in previous work [97]) supplemented with Vancomycin (Vc; 7.5 μg/μl), Chloramphenicol (Cm; 0.5μg/μl) and Erythromycin (Ery; 14.6μg/μl), in addition to Gifu Anaerobic Broth, Modified agar plates (mGAM; HiMedia M2079). SK (Vc, Cm and Ery) and mGAM agar plates (1.5% agar, Fisher Scientific, 10572775) were prepared. For plates supplemented with Vc, Cm and Ery, antibiotics were added during the plates preparation. All plates were transferred to the anaerobic chamber under anaerobic conditions (5% H2, 10% CO2 and 85% N2) to be pre-reduced for 24h before plating. Frozen saliva and faecal samples were thawed in the anaerobic chamber and serial dilution were realised, where one hundred microliters of diluted samples were dispensed and spread on the plates. The plates were incubated for up to 4 days at 37°C in the anaerobic chamber for colony growth.

After each day, strain imaging and colony picking was performed for further isolation. The colony picking was based on the identification of morphologically unique colonies including area perimeter, circularity, convexity, color and consistency. Colonies were stricken on the equivalent media from which they were picked and incubated at 37°C in the anaerobic chamber for colony growth.

### Bacterial strains and growth conditions

**Table S1** lists the bacterial strains used in this study. Strains were grown at 37°C in an anaerobic chamber (Whitley M45, Meintrup DWS Laborgeräte GmbH) at an atmosphere of 5% H_2_, 5% CO_2_, and 90% N_2_. MS082 was cultivated on mGAM agar plates (modified Gifu Anaerobe Medium, Himedia). Strains MS055, MS061, MS072, MS097, MS107 and MS164 were cultivated in SK broth or on agar plates (Difco^TM^ tryptone 10 g/l, yeast extract 10 g/l, NaCl 2 g/l, Na_2_HPO_4_ 0.4 g/l) [35].

### Growth curve characteristics of *Veillonella* strains

The strains MS055 and MS061 were grown in SK broth as detailed under growth conditions, and in SK medium supplemented with 50 mM DL-lactate (Sigma-Aldrich), or 50 mM potassium nitrate (Sigma-Aldrich), respectively. In brief, overnight cultures of MS055 and MS061 in SK medium were diluted into new medium for log-phase growth and used to inoculate the main culture of 6 ml medium (start OD_600_ 0.05) of SK medium only, and SK medium with 50 mM lactate or nitrate, respectively. Growth of strains was monitored every 2 hours using a spectrophotometer (CO8000, Biowave). The results represent three independent experiments and are presented as means with standard errors of the means.

### Gene neighbourhood visualisation from *de novo* assemblies

For each gene family of interest, we retrieved all assembled contigs containing this gene. Then we aligned the contigs centred to the gene of interest taking strand information into account. The gene families in the same relative position to the centre gene were retrieved and the most prevalent gene family at each relative position was visualised including the number of observations of that gene.

### Identification of *prtC* homologs in metagenomic species pangenomes

Each candidate oral-gut translocator species was first matched to their corresponding MSPs based on gtdb-tk taxonomic annotations of the assembled MSPs and linear correlations of the respective MetaPhlAn- and MSP-abundances (for further details see section “Integration of reference- and assembly-based results”). We were able to assign MPSs for seven out of nine oral-gut translocators. Functional group (FG) annotations for each gene family in the MSPs were extracted and 288 FGs were found to be commonly shared among MSPs associated with oral-gut translocation (>80% prevalence). Next, we excluded functional groups that represent functions commonly found in the gut, by comparing these 288 FGs to those present in the core genomes of prevalent gut commensals. For this comparison, we selected MSPs with a mean abundance greater than 5% across all healthy faecal samples, including *Prevotella copri*, *Bacteroides uniformis*, *and Faecalibacterium prausnitzii*. FGs were filtered if they were also present in these common healthy gut microbes and this filtering step reduced the number of remaining FGs to 52, which may represent bacterial functions that are essential for oral-gut translocation and/or result in aberrant host-microbial interactions in disease. Cumulative faecal abundances for these 52 FGs were calculated by summing over all matched gene families in oral-gut translocators. Subsequently, we further prioritised these FGs according to the correlations between their faecal abundance and plasma FABP2 levels. For this, the pcor.test function in R (rank-based Spearman) was used to calculate partial correlations with age, gender, and antibiotic usage included as covariates. P-values were corrected for multiple testing with Benjamini Hochberg.

### Predicting ACLD based on faecal *prtC* gene abundance

First, the cumulative abundance of *prtC* genes encoded by oral-gut translocators was inferred for each faecal sample and represented as a single value with the detection limit set to 0.5 RPKM. These values were then used to predict disease status: any sample with a value above this threshold was predicted as “ACLD” and samples below this value as “control”. The confusion matrix, specificity and sensitivity is reported in **Table 2**. The corresponding Receiver Operating Characteristic (ROC) and Precision-Recall (PR) curves were generated using the cumulative *prtC* abundance as the predictor and actual disease status (based on the clinical metadata) as the ground truth (1 for ACLD, 0 for controls)

### Verification of the *prtC* signals in CLD-Sole2021 cohort

The cohort by Solé et al. [11] [CLD-Sole2021] contained metagenomic data with single-end reads based on ion proton sequencing. Due to the shorter read length and lower read quality we did not assemble this dataset but employed a mapping-based strategy (bowtie2) against the gene catalogue generated from our cohort. RPKM values are calculated from reads mapped to the *prtC* genes.

### Structure comparisons between *prtC* genes and a known bacterial collagenase

We applied blastx to search the UniProt database using gene sequences of the *prtC* genes identified in our cohort (**Table 2**). The best hit for each query sequence was taken for further analysis, where the predicted protein structure was downloaded as .pdb format and was aligned against the structure of a well-characterised bacterial collagenase (P33437). In the cases where multiple queries matched to the same protein from the UniProt database, we only took one matched protein for the downstream analysis. For example, the *prtC* genes from *V. parvula*, *V. atypcia* and *V. dispar* were all mapped to *Veillonella sp. HPA0037*, therefore we used the predicted *prtC* structure from *Veillonella sp. HPA0037* for the alignment to represent these three *prtC* genes. To quantify the structural similarity, the root-mean-square-deviation (RMSD) was calculated for each structure alignment.

### Experimental procedures in animals

*In vivo* experimental procedures were performed in 10-12-week-old male C57/Bl6 mice at the Disease Modelling Unit (University of East Anglia, UK). Experiments were approved by the Animal Welfare and Ethical Review Body (AWERB, University of East Anglia, Norwich, UK) and the UK Home Office (Project licence to Beraza: PP9417531). All procedures were carried out following the guidelines of the National Academy of Sciences (National Institutes of Health, publication 86-23, revised 1985) and were performed within the provisions of the Animals (Scientific Procedures) Act 1986 (ASPA).

### Induction of hepatic fibrosis *in vivo*

Mice were treated with CCl_4_ (1ml/kg) that was administered via IP twice per week for a total of 6 weeks. Two weeks after the initiation of CCl_4_ treatment, one group of mice received 200ul of a bacterial cocktail (10^9^ CFU/ml) composed of *Veillonella parvula* (isolates MS061, MS107 and MS164), *Veillonella dispar* (isolate MS072) and *Streptococcus parasanguinis* (isolate MS082) by oral gavage. The strain cocktail was administered for three consecutive days each week for 4 weeks (week 3 to week 6 of treatment). All mice were sacrificed two days after the last administration of CCl_4_ (**Figure 4a**).

### Quantification of fibrosis severity in mice

Liver tissues were fixed in 10% neutral buffered formalin (Sigma–HT501128-4L), embedded in paraffin and sectioned. Liver sections were dewaxed, hydrated and stained with Sirius red to stain collagen and detect fibrosis. Slides were imaged on a BX53 upright microscope (Olympus) with an Olympus DP74 colour camera and a pT100 LED transmitted light source (CoolLED). For quantification of the deposition of collagen in the liver, a total of 6-10 fields of view per sample were imaged and analysed using open-source FIJI software [98] as described previously [99]. Fibrosis is represented as the percentage of stained area relative to total area per field.

### Quantification of gut barrier dysfunction in mice

Frozen faecal material from the large intestine was weighed and diluted in a 1:15 ratio (weight:volume) using extraction buffer (50 mM Tris HCl, 150 mM NaCl, 0.1% SDS, 2 mM EDTA (pH 8.0)). Samples were homogenised 3 times at 6 m/s for 40 seconds and homogenates were centrifuged at 12,000 rpm at 4C for 10 minutes. Supernatants were collected and assessed for Albumin levels. Albumin levels were quantified using the DY1455 human albumin Duoset enzyme-linked immunosorbent assay (ELISA) from R&D Systems (Minneapolis, MN, USA) according to manufacturer instructions. Results were quantified using a FluoStar Optima plate reader. The mouse gastrointestinal tract was dissected into anatomically defined regions, and the terminal portion of the ileum and the colon were fixed and embedded in paraffin for immunohistological analysis: slides were mounted with a DAPI-mounting solution (Vector Laboratories – H-1200) to stain cell nuclei. Fluorescent microscopic imaging was performed using an AxioImager M2 (Zeiss) with the AxioCam mRM monochrome camera and standard light source and filter sets supplied.

### Preparation of faecal water

Frozen faecal samples from AD (n = 10) and healthy (n = 10) individuals were thawed on ice. A 0.3g aliquot from each sample was suspended in 2 mL of 1× phosphate-buffered saline (PBS) and homogenized thoroughly. The homogenates were centrifuged at 5,000 rpm for 30 minutes at 4 °C. The resulting supernatants were collected and subjected to two additional centrifugation steps at 5,000 rpm for 20 minutes and 10 minutes, respectively, both at 4 °C. Supernatants were collected and kept on ice after each step for downstream analysis.

### *E.coli* BL21 (DE3) pET29b(+)/*prtC* transformation

The *prtC* gene sequence from *Veillonella parvula* (isolate MS055) was synthetised and cloned in pEt29b(+) by TwistBioscience. Competent *E.coli* BL-21 was transformed by heat shock with the pET-29b(+)/*prtC* vector and plated on Luria-Broth agar media supplemented with Kanamycin (25ug/mL) and incubated 24h at 37°C.

### Recombinant protein expression and preparation

*E. coli BL21* (DE3) and BL21 (DE3) harboring the pET-29b(+)/*prtC* plasmid were cultured in 100 mL Luria Broth at 25 °C with shaking at 150 rpm until reaching an optical density at 600 nm (OD₆₀₀) of 0.8. Protein expression in BL21 (DE3) pET-29b(+)/*prtC* was induced with M isopropyl β-D-1-thiogalactopyranoside (IPTG), followed by incubation at 25 °C for 18 hours. Induced and non-induced cultures were centrifuged at 5,000 rpm for 20 minutes at 4 °C. The resulting supernatants were transferred to Amicon® Ultra Centrifugal Filter tubes and centrifuged again at 5,000 rpm for 10 minutes at 4 °C.

### Collagenase activity in faecal water / Collagen degradation Assay

Collagen degradation was assessed using the EnzChek Gelatinase/Collagenase Assay Kit (Thermo Fisher Scientific). DQ-collagen I was added to concentrated supernatants, a media-only control, and a positive control at a final concentration of 100 µg/mL. Samples were incubated overnight at 37 °C. Fluorescence was measured at an excitation wavelength of 495 nm and emission at 515 nm. The fluorescence intensity was directly proportional to the extent of collagen degradation.

## References

1. Cheemerla, S. and M. Balakrishnan, Global epidemiology of chronic liver disease. Clinical Liver Disease, 2021. 17(5): p. 365.

2. Asrani, S.K., et al., Burden of liver diseases in the world. Journal of hepatology, 2019. 70(1): p. 151–171.

3. Ginès, P., et al., Liver cirrhosis. The Lancet, 2021. 398(10308): p. 1359–1376.

4. Borthwick, L. and F. Oakley, Editorial overview: fibrosis. 2019. p. vi–vii.

5. D’Amico, G., M. Bernardi, and P. Angeli, Towards a new definition of decompensated cirrhosis. Journal of hepatology, 2021.

6. Moreau, R., et al., Acute-on-chronic liver failure: a distinct clinical syndrome. Journal of hepatology, 2021. 75: p. S27–S35.

7. Teerasarntipan, T., et al., Validation of prognostic scores predicting mortality in acute liver decompensation or acute-on-chronic liver failure: A Thailand multicenter study. Plos one, 2022. 17(11): p. e0277959.

8. Yoshiji, H., et al., Evidence-based clinical practice guidelines for liver cirrhosis 2020. Journal of Gastroenterology, 2021. 56(7): p. 593–619.

9. Singal, A.K. and P. Mathurin, Diagnosis and treatment of alcohol-associated liver disease: a review. Jama, 2021. 326(2): p. 165–176.

10. Scaglione, S., et al., The epidemiology of cirrhosis in the United States. Journal of clinical gastroenterology, 2015. 49(8): p. 690–696.

11. Solé, C., et al., Alterations in gut microbiome in cirrhosis as assessed by quantitative metagenomics: relationship with acute-on-chronic liver failure and prognosis. Gastroenterology, 2021. 160(1): p. 206–218. e13.

12. Qin, N., et al., Alterations of the human gut microbiome in liver cirrhosis. Nature, 2014. 513(7516): p. 59–64.

13. Chen, Y., et al., Characterization of fecal microbial communities in patients with liver cirrhosis. Hepatology, 2011. 54(2): p. 562–572.

14. Acharya, C. and J.S. Bajaj, Altered microbiome in patients with cirrhosis and complications. Clinical Gastroenterology and Hepatology, 2019. 17(2): p. 307–321.

15. Bajaj, J.S., et al., Gut microbiota alterations can predict hospitalizations in cirrhosis independent of diabetes mellitus. Scientific reports, 2015. 5(1): p. 18559.

16. Leung, H., et al., Impaired flux of bile acids from the liver to the gut reveals microbiome-immune interactions associated with liver damage. npj Biofilms and Microbiomes, 2023. 9(1): p. 35.

17. Cabrera-Rubio, R., et al., Cholestasis induced by bile duct ligation promotes changes in the intestinal microbiome in mice. Scientific reports, 2019. 9(1): p. 12324.

18. Avila, M.A., et al., Recent advances in alcohol-related liver disease (ALD): summary of a Gut round table meeting. Gut, 2020. 69(4): p. 764–780.

19. Leung, H., et al., Risk assessment with gut microbiome and metabolite markers in NAFLD development. Science translational medicine, 2022. 14(648): p. eabk0855.

20. Oh, T.G., et al., A universal gut-microbiome-derived signature predicts cirrhosis. Cell metabolism, 2020. 32(5): p. 878–888. e6.

21. Qiu, Q., et al., Metagenomic analysis reveals the distribution of antibiotic resistance genes in a large-scale population of healthy individuals and patients with varied diseases. Frontiers in Molecular Biosciences, 2020. 7: p. 590018.

22. Ali, R.O., et al., Longitudinal multi-omics analyses of the gut–liver axis reveals metabolic dysregulation in hepatitis C infection and cirrhosis. Nature Microbiology, 2022: p. 1–16.

23. Acharya, C., S.E. Sahingur, and J.S. Bajaj, Microbiota, cirrhosis, and the emerging oral-gut-liver axis. JCI insight, 2017. 2(19).

24. Schirmer, M., et al., Linking microbial genes to plasma and stool metabolites uncovers host-microbial interactions underlying ulcerative colitis disease course. Cell Host & Microbe, 2024. 32(2): p. 209–226. e7.

25. Zeller, G., et al., Potential of fecal microbiota for early-stage detection of colorectal cancer. Molecular systems biology, 2014. 10(11): p. 766.

26. Thomas, A.M., et al., Metagenomic analysis of colorectal cancer datasets identifies cross-cohort microbial diagnostic signatures and a link with choline degradation. Nature medicine, 2019. 25(4): p. 667–678.

27. Schmidt, T.S., et al., Extensive transmission of microbes along the gastrointestinal tract. Elife, 2019. 8: p. e42693.

28. Vatanen, T., et al., The human gut microbiome in early-onset type 1 diabetes from the TEDDY study. Nature, 2018. 562(7728): p. 589–594.

29. Chen, B.-Y., et al., Roles of oral microbiota and oral-gut microbial transmission in hypertension. Journal of Advanced Research, 2023. 43: p. 147–161.

30. Valles-Colomer, M., et al., The person-to-person transmission landscape of the gut and oral microbiomes. Nature, 2023. 614(7946): p. 125–135.

31. Jin, S., D. Wetzel, and M. Schirmer, Deciphering mechanisms and implications of bacterial translocation in human health and disease. Current Opinion in Microbiology, 2022. 67: p. 102147.

32. Patel, V.C., et al., Rifaximin-α reduces gut-derived inflammation and mucin degradation in cirrhosis and encephalopathy: RIFSYS randomised controlled trial. Journal of hepatology, 2022. 76(2): p. 332–342.

33. Liao, C., et al., Oral bacteria relative abundance in faeces increases due to gut microbiota depletion and is linked with patient outcomes. Nature microbiology, 2024: p. 1–11.

34. Lloyd-Price, J., et al., Multi-omics of the gut microbial ecosystem in inflammatory bowel diseases. Nature, 2019. 569(7758): p. 655–662.

35. Rojas-Tapias, D.F., et al., Inflammation-associated nitrate facilitates ectopic colonization of oral bacterium Veillonella parvula in the intestine. Nature Microbiology, 2022. 7(10): p. 1673–1685.

36. Winter, S.E., et al., Host-derived nitrate boosts growth of E. coli in the inflamed gut. science, 2013. 339(6120): p. 708–711.

37. Beghini, F., et al., Integrating taxonomic, functional, and strain-level profiling of diverse microbial communities with bioBakery 3. Elife, 2021. 10: p. e65088.

38. Truong, D.T., et al., Microbial strain-level population structure and genetic diversity from metagenomes. Genome research, 2017. 27(4): p. 626–638.

39. Scholz, M., et al., Strain-level microbial epidemiology and population genomics from shotgun metagenomics. Nature methods, 2016. 13(5): p. 435–438.

40. Peng, Y., X. Qi, and X. Guo, Child–Pugh versus MELD score for the assessment of prognosis in liver cirrhosis: a systematic review and meta-analysis of observational studies. Medicine, 2016. 95(8).

41. Storch, J. and B. Corsico, The emerging functions and mechanisms of mammalian fatty acid–binding proteins. Annual review of nutrition, 2008. 28.

42. Gajda, A.M. and J. Storch, Enterocyte fatty acid-binding proteins (FABPs): different functions of liver and intestinal FABPs in the intestine. Prostaglandins, Leukotrienes and Essential Fatty Acids, 2015. 93: p. 9–16.

43. Levy, E., et al., Localization, function and regulation of the two intestinal fatty acid-binding protein types. Histochemistry and cell biology, 2009. 132(3): p. 351–367.

44. Riva, A., et al., Faecal cytokine profiling as a marker of intestinal inflammation in acutely decompensated cirrhosis. JHEP Reports, 2020. 2(6): p. 100151.

45. Bajaj, J.S., P.S. Kamath, and K.R. Reddy, The evolving challenge of infections in cirrhosis. New England Journal of Medicine, 2021. 384(24): p. 2317–2330.

46. Tripathi, A., et al., The gut–liver axis and the intersection with the microbiome. Nature reviews Gastroenterology & hepatology, 2018. 15(7): p. 397–411.

47. Llorente, C. and B. Schnabl, The gut microbiota and liver disease. Cellular and molecular gastroenterology and hepatology, 2015. 1(3): p. 275–284.

48. Keshavarzian, A., et al., Leaky gut in alcoholic cirrhosis: a possible mechanism for alcohol-induced liver damage. The American journal of gastroenterology, 1999. 94(1): p. 200–207.

49. Kato, T., N. Takahashi, and H.K. Kuramitsu, Sequence analysis and characterization of the Porphyromonas gingivalis prtC gene, which expresses a novel collagenase activity. Journal of bacteriology, 1992. 174(12): p. 3889–3895.

50. Wu, Y.m., et al., Effect of Porphyromonas gingivalis PrtC on cytokine expression in ECV304 endothelial cells and its level in subgingival plaques from patients with chronic periodontitis 1. Acta pharmacologica sinica, 2007. 28(7): p. 1015–1023.

51. Bonnans, C., J. Chou, and Z. Werb, Remodelling the extracellular matrix in development and disease. Nature reviews Molecular cell biology, 2014. 15(12): p. 786–801.

52. Albuquerque-Souza, E. and S.E. Sahingur, Periodontitis, chronic liver diseases, and the emerging oral-gut-liver axis. Periodontology 2000, 2022. 89(1): p. 125–141.

53. Schirmer, M., et al., Linking the human gut microbiome to inflammatory cytokine production capacity. Cell, 2016. 167(4): p. 1125–1136. e8.

54. Tigchelaar, E.F., et al., Cohort profile: LifeLines DEEP, a prospective, general population cohort study in the northern Netherlands: study design and baseline characteristics. BMJ open, 2015. 5(8): p. e006772.

55. Zhernakova, A., et al., Population-based metagenomics analysis reveals markers for gut microbiome composition and diversity. Science, 2016. 352(6285): p. 565–569.

56. Hu, J., et al., Enteric dysbiosis-linked gut barrier disruption triggers early renal injury induced by chronic high salt feeding in mice. Experimental & molecular medicine, 2017. 49(8): p. e370–e370.

57. Wang, L., et al., Methods to determine intestinal permeability and bacterial translocation during liver disease. Journal of immunological methods, 2015. 421: p. 44–53.

58. Zuo, L., W.-T. Kuo, and J.R. Turner, Tight junctions as targets and effectors of mucosal immune homeostasis. Cellular and molecular gastroenterology and hepatology, 2020. 10(2): p. 327–340.

59. Shi, D., et al., Administration of Lactobacillus salivarius LI01 or Pediococcus pentosaceus LI05 prevents CCl4-induced liver cirrhosis by protecting the intestinal barrier in rats. Scientific reports, 2017. 7(1): p. 6927.

60. Rieder, F., et al., Intestinal fibrosis and liver fibrosis: consequences of chronic inflammation or independent pathophysiology? Inflammatory intestinal diseases, 2016. 1(1): p. 41–49.

61. Kessoku, T., et al., The role of leaky gut in nonalcoholic fatty liver disease: a novel therapeutic target. International Journal of Molecular Sciences, 2021. 22(15): p. 8161.

62. Fukui, H., Gut-liver axis in liver cirrhosis: How to manage leaky gut and endotoxemia. World journal of hepatology, 2015. 7(3): p. 425.

63. Nishimura, N., et al., Intestinal permeability is a mechanical rheostat in the pathogenesis of liver cirrhosis. International Journal of Molecular Sciences, 2021. 22(13): p. 6921.

64. Bedi, G.S. and T. Williams, Purification and characterization of a collagen-degrading protease from Porphyromonas gingivalis. Journal of Biological Chemistry, 1994. 269(1): p. 599–606.

65. Wang, X., et al., Gut–liver translocation of pathogen Klebsiella pneumoniae promotes hepatocellular carcinoma in mice. Nature Microbiology, 2025. 10(1): p. 169–184.

66. Soresi, M., et al., Non invasive tools for the diagnosis of liver cirrhosis. World journal of gastroenterology: WJG, 2014. 20(48): p. 18131.

67. Denzer, U.W. and S. Lüth, Non-invasive diagnosis and monitoring of liver fibrosis and cirrhosis. Best practice & research Clinical gastroenterology, 2009. 23(3): p. 453–460.

68. Speca, S., et al., Cellular and molecular mechanisms of intestinal fibrosis. World journal of gastroenterology: WJG, 2012. 18(28): p. 3635.

69. Wu, X., et al., Cellular and molecular mechanisms of intestinal fibrosis. Gut and Liver, 2023. 17(3): p. 360.

70. Maslennikov, R., et al., Gut microbiota and bacterial translocation in the pathogenesis of liver fibrosis. International journal of molecular sciences, 2023. 24(22): p. 16502.

71. Fukui, H., Leaky gut and gut-liver axis in liver cirrhosis: clinical studies update. Gut and liver, 2021. 15(5): p. 666.

72. Trebicka, J., et al., Utilizing the gut microbiome in decompensated cirrhosis and acute-on-chronic liver failure. Nature reviews Gastroenterology & hepatology, 2021. 18(3): p. 167–180.

73. Furuichi, M., et al., Commensal consortia decolonize Enterobacteriaceae via ecological control. Nature, 2024. 633(8031): p. 878–886.

74. Lee, S., et al., Oral-gut microbiome interactions in advanced cirrhosis: characterisation of pathogenic enterotypes and salivatypes, virulence factors and antimicrobial resistance. Journal of Hepatology, 2024.

75. Angeli, P., et al., EASL Clinical Practice Guidelines for the management of patients with decompensated cirrhosis. Journal of hepatology, 2018. 69(2): p. 406–460.

76. Moreau, R., et al., Acute-on-chronic liver failure is a distinct syndrome that develops in patients with acute decompensation of cirrhosis. Gastroenterology, 2013. 144(7): p. 1426–1437. e9.

77. Arroyo, V., et al., Acute-on-chronic liver failure in cirrhosis. Nature reviews Disease primers, 2016. 2(1): p. 1–18.

78. Pugh, R., et al., Transection of the oesophagus for bleeding oesophageal varices. British journal of surgery, 1973. 60(8): p. 646–649.

79. Abramowicz, M., Hyponatremia and mortality among patients waiting for liver transplantation. The New England journal of medicine, 2008. 359(24): p. 2615.

80. Singer, M., et al., The third international consensus definitions for sepsis and septic shock (Sepsis-3). Jama, 2016. 315(8): p. 801–810.

81. Dore, J., et al., Standard Operating Procedure for Fecal Samples DNA Extraction. Protocol Q IHMS Consortium. IHMS_SOP, 2015. 6: p. V2.

82. Costea, P.I., et al., Towards standards for human fecal sample processing in metagenomic studies. Nature biotechnology, 2017. 35(11): p. 1069–1076.

83. Suzuki, M.T., L.T. Taylor, and E.F. DeLong, Quantitative analysis of small-subunit rRNA genes in mixed microbial populations via 5′-nuclease assays. Applied and environmental microbiology, 2000. 66(11): p. 4605–4614.

84. Harju, S., H. Fedosyuk, and K.R. Peterson, Rapid isolation of yeast genomic DNA: Bust n’ Grab. BMC Biotechnology, 2004. 4(1): p. 8.

85. Yao, B., et al., Quantification and characterization of mouse and human tissue-resident microbiota by qPCR and 16S sequencing. STAR protocols, 2022. 3(4): p. 101765.

86. Andrews, S., et al., Trim Galore. Trim Galore, 2015.

87. Li, D., et al., MEGAHIT: an ultra-fast single-node solution for large and complex metagenomics assembly via succinct de Bruijn graph. Bioinformatics, 2015. 31(10): p. 1674–1676.

88. Hyatt, D., et al., Gene and translation initiation site prediction in metagenomic sequences. Bioinformatics, 2012. 28(17): p. 2223–2230.

89. Hyatt, D., et al., Prodigal: prokaryotic gene recognition and translation initiation site identification. BMC bioinformatics, 2010. 11(1): p. 1–11.

90. Li, W. and A. Godzik, Cd-hit: a fast program for clustering and comparing large sets of protein or nucleotide sequences. Bioinformatics, 2006. 22(13): p. 1658–1659.

91. Fu, L., et al., CD-HIT: accelerated for clustering the next-generation sequencing data. Bioinformatics, 2012. 28(23): p. 3150–3152.

92. Aroney, S.T., et al., CoverM: read alignment statistics for metagenomics. Bioinformatics, 2025. 41(4): p. btaf147.

93. Quevillon, E., et al., InterProScan: protein domains identifier. Nucleic acids research, 2005. 33(suppl_2): p. W116–W120.

94. Jones, P., et al., InterProScan 5: genome-scale protein function classification. Bioinformatics, 2014. 30(9): p. 1236–1240.

95. Plaza Oñate, F., et al., MSPminer: abundance-based reconstitution of microbial pan-genomes from shotgun metagenomic data. Bioinformatics, 2019. 35(9): p. 1544–1552.

96. Chaumeil, P.-A., et al., GTDB-Tk v2: memory friendly classification with the genome taxonomy database. Bioinformatics, 2022. 38(23): p. 5315–5316.

97. Béchon, N., et al., Autotransporters drive biofilm formation and autoaggregation in the diderm firmicute Veillonella parvula. Journal of Bacteriology, 2020. 202(21): p. 10.1128/jb.00461-20.

98. Schindelin, J., et al., Fiji: an open-source platform for biological-image analysis. Nature methods, 2012. 9(7): p. 676–682.

99. Moreno-Gonzalez, M., et al., Regulation of intestinal senescence during cholestatic liver disease modulates barrier function and liver disease progression. JHEP Reports, 2024. 6(10): p. 101159.

