## Supplementary figures for "Microbial oral-gut translocation in advanced chronic liver disease is linked to exacerbation of intestinal barrier dysfunction and hepatic fibrosis"

**Figure S1**

a

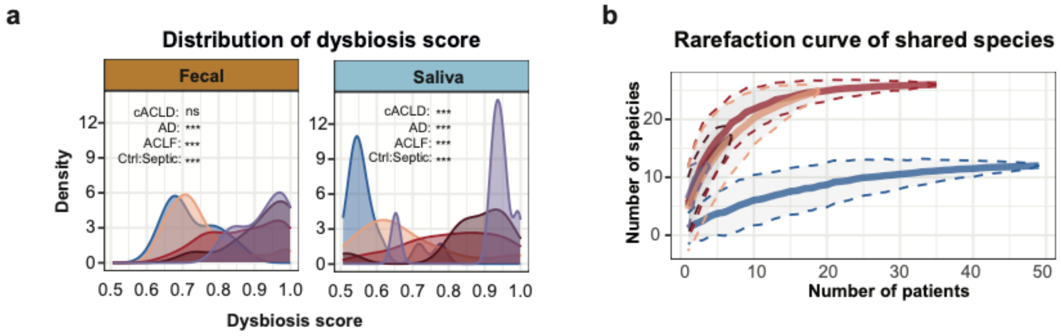


c

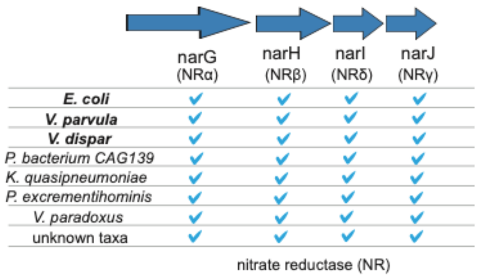


d

Candidate oral-gut translocators

| Species | Gene profiling | SNV profiling |
|---|---|---|
| *Haemophilus parainfluenzae* | Yes | No |
| *Megasphaera micronuciformis* | No | Yes |
| *Rothia mucilaginosa* | Yes | Yes |
| *Streptococcus parasanguinis* | Yes | Yes |
| *Streptococcus salivarius* | Yes | Yes |
| *Veillonella atypica* | Yes | Yes |
| *Veillonella dispar* | Yes | No |
| *Veillonella infantium* | No | No |
| *Veillonella parvula* | Yes | Yes |

e

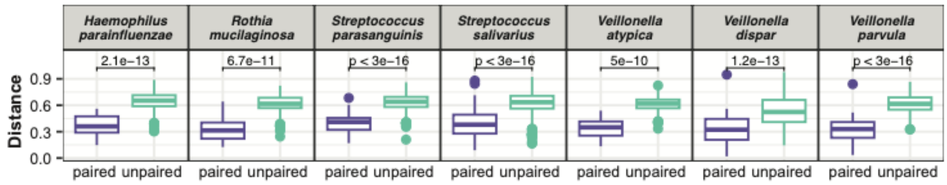


f

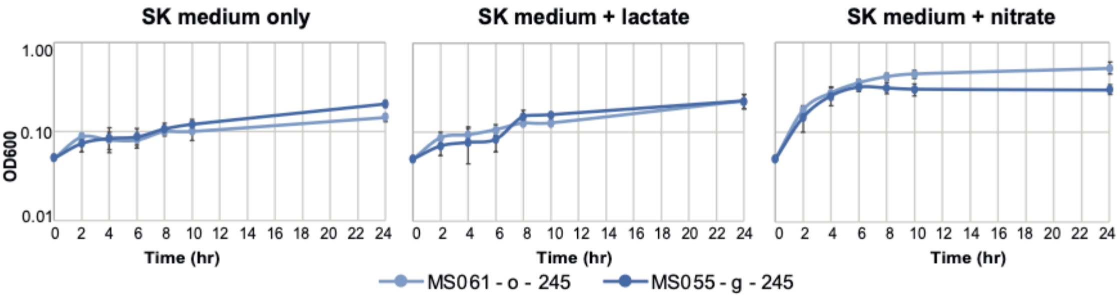


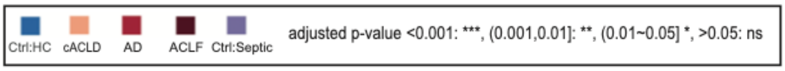

**Figure S1.** **(a)** Dysbiosis score distribution of faecal (left) and saliva (right) microbiome (Wilcoxon). **(b)** Rarefaction curves showing number of shared species against number of patients for each disease group (95% confidence interval). **(c)** Presence of the nitrate reductase (nar) operon in metagenomic assembled pangenomes (MSPs). The presence of four key genes of the nar operon were examined. Species with all narGHJI genes (>30% identity >80% coverage) are shown and species commonly shared between faecal and saliva samples are marked in bold. **(d)** Summary table of identified oral-gut translocators and their available strain profiles based on gene content and single nucleotide variation (SNV) analysis. **(e)** Comparison of gene content similarity between strains from paired versus unpaired faecal and saliva samples. Y-axis reflects binary distance estimates based on gene presence/absence in the respective pangenome (PanPhlAn). **(f)** Next-to-identical *V. parvula* strains from paired faecal and saliva samples show similar growth characteristics under various conditions (SK medium +/- lactate or nitrate).

### Figure S2

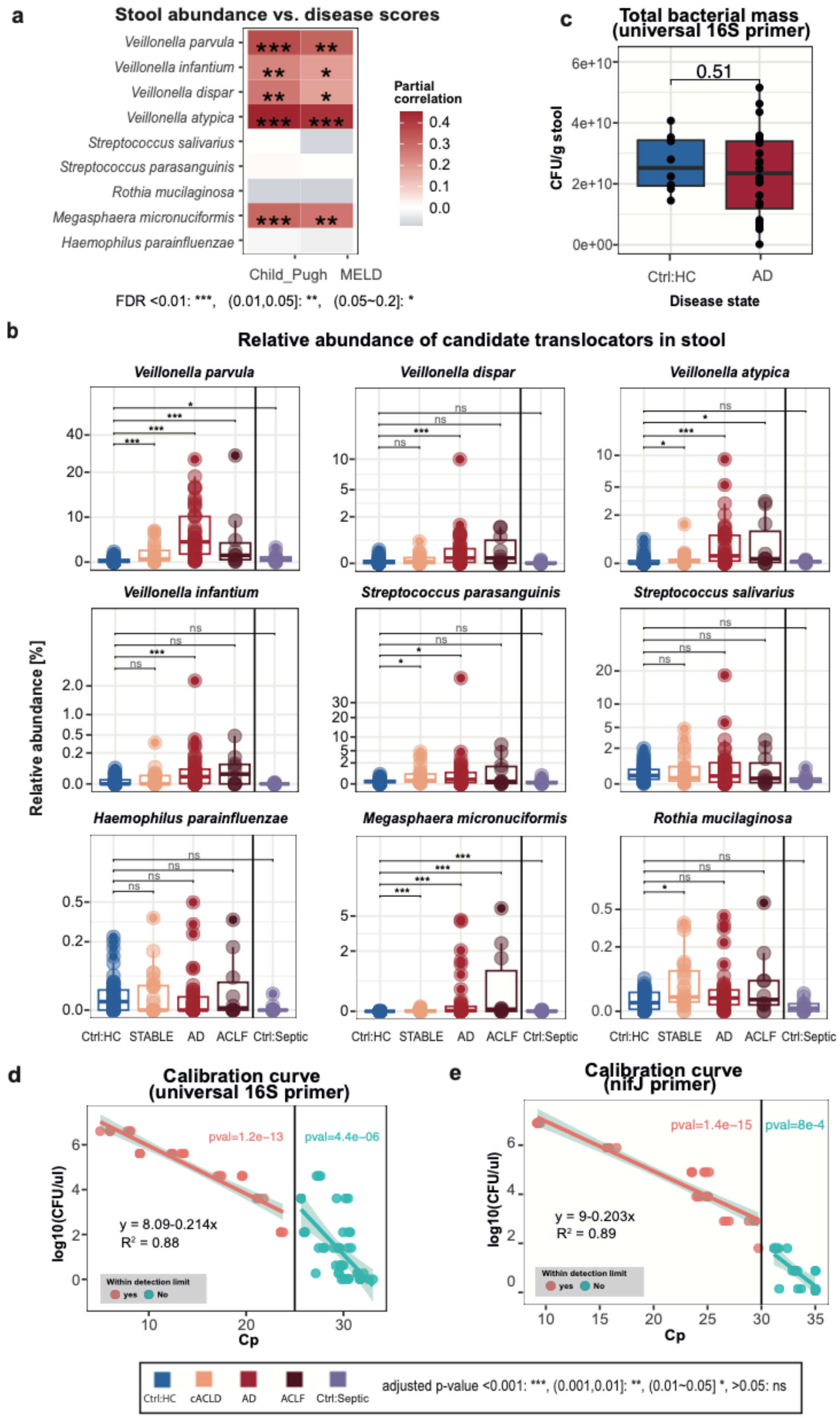
a
Stool abundance vs. disease scores
Veillonella parvula
Veillonella infantium
Veillonella dispar
Veillonella atypica
Streptococcus salivarius
Streptococcus parasanguinis
Rothia mucilaginosa
Megasphaera micronuciformis
Haemophilus parainfluenzae
Child_Pugh
MELD
Partial correlation
FDR <0.01: ***, (0.01,0.05]: **, (0.05~0.2]: *
c
Total bacterial mass (universal 16S primer)
CFU/g stool
0.51
Ctrl:HC
AD
Disease state
b
Relative abundance of candidate translocators in stool
Relative abundance [%]
Veillonella parvula
Veillonella dispar
Veillonella atypica
Veillonella infantium
Streptococcus parasanguinis
Streptococcus salivarius
Haemophilus parainfluenzae
Megasphaera micronuciformis
Rothia mucilaginosa
Ctrl:HC STABLE AD ACLF Ctrl:Septic
d
Calibration curve (universal 16S primer)
log10(CFU/ul)
pval=1.2e−13
pval=4.4e−06
y = 8.09-0.214x
R² = 0.88
Within detection limit
yes No
Cp
e
Calibration curve (nifJ primer)
log10(CFU/ul)
pval=1.4e−15
pval=8e−4
y = 9-0.203x
R² = 0.89
Within detection limit
yes No
Cp
Ctrl:HC cACLD AD ACLF Ctrl:Septic
adjusted p-value <0.001: ***, (0.001,0.01]: **, (0.01~0.05] *, >0.05: ns

**Figure S2 (a)** Correlation of faecal abundance of translocating species with Child-Pugh and MELD scores (partial correlation, Spearman, corrected for age, gender antibiotic usage, p-values were adjusted with BH, adjusted p-value <0.01: ***, (0.01,0.05]: **, (0.05~0.2]: *) **(b)** Relative abundance of each oral-gut translocator in faeces per disease group (one-sided Wilcoxon, assessing whether total faecal abundance of translocators is increased in ACLD group, p-value adjusted with BH). **(c)** Total bacterial load estimation by qPCR showed comparable levels across disease groups (samples from 25 acutely decompensated ACLD (AD) and 8 healthy controls (Ctrl:HC) with on average (median) $23\times10^9$ and $25\times10^9$ colony-forming units (CFUs), respectively (Wilcoxon test). **(d)** Calibration curve based on universal 16S rRNA primer. Y-axis indicates CFU/ul for *Escherichia coli* and the x-axis shows the measured Cp value. Red points are within the detection limits (25 Cp) and the equation shows the fitted linear regression, p-value=$1.2e^{-13}$. **(e)** Analogous calibration curve for the *nifJ* primer targeting *V. parvula* {Rojas-Tapias, 2022 #435}. (Detection limits = 30 Cp and fitted linear regression with p-value=$1.4e^{-15}$.)

#### Figure S3

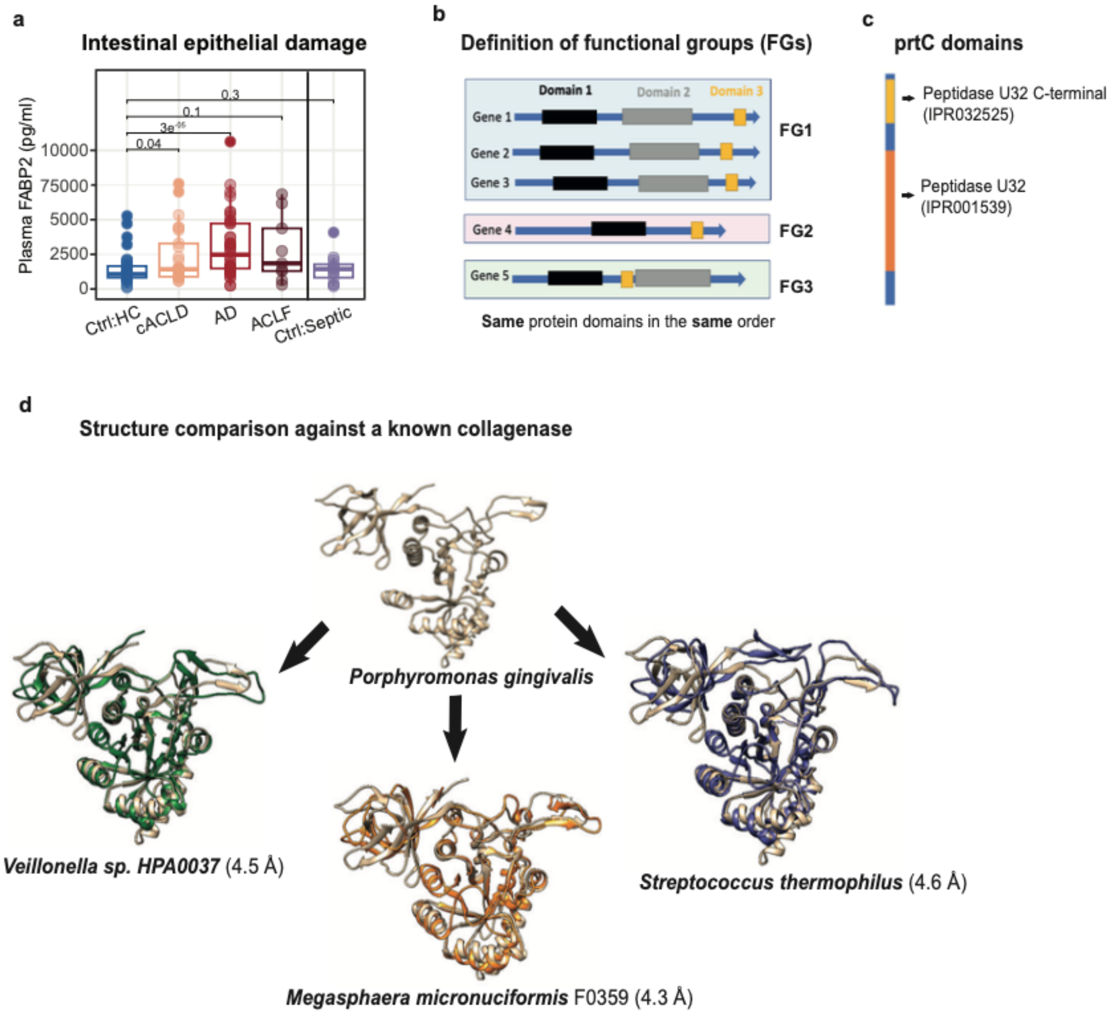


**Figure S3 (a)** Fatty acid binding protein 2 (FABP2) levels (y-axis) were compared across different disease groups (Wilcoxon, alternative hypothesis: FABP2 reduced in Ctrl:HC, p-value adjusted with BH). **(b)** Schematic for the definition of functional groups (FGs), where protein encoding genes are grouped if they share the same functional domains in the same order. **(c)** Functional domain organisation of the prtC gene. **(d)** Structural comparison between *prtC* genes identified from ACLD patients and a well-characterised *prtC* gene from *Porphyromonas gingivitis* (P33437). The closest homologs in UniProt were identified using blastx (**Table 2**), structures were predicted by AlphaFold2 and then aligned against the *P. gingivalis prtC* gene. Structure differences are measured by root-mean-square-deviation (RMSD) in angstrom (Å)**.**

#### Figure S4

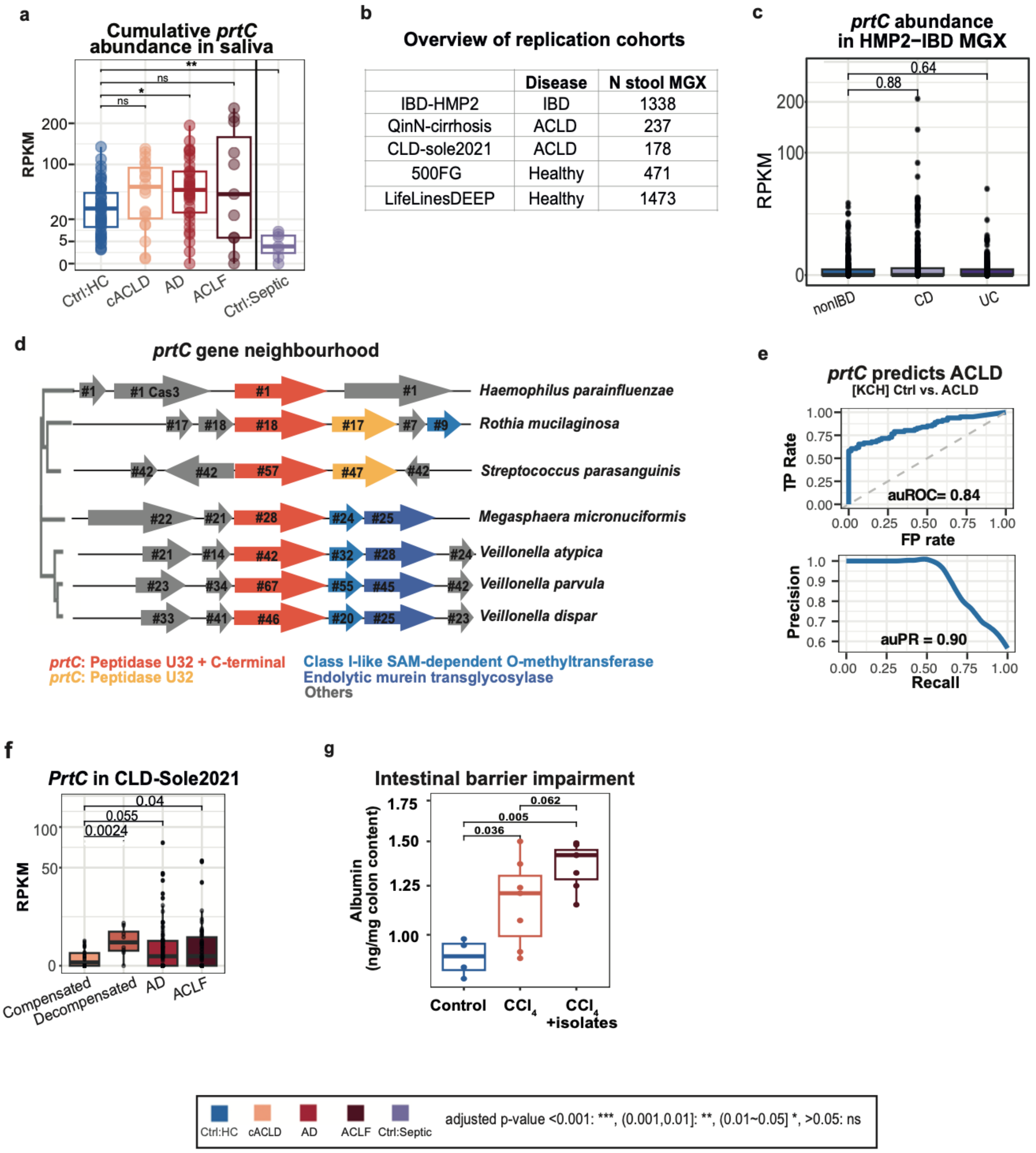


| | Disease | N stool MGX |
|---|---|---|
| IBD-HMP2 | IBD | 1338 |
| QinN-cirrhosis | ACLD | 237 |
| CLD-sole2021 | ACLD | 178 |
| 500FG | Healthy | 471 |
| LifeLinesDEEP | Healthy | 1473 |

**Figure S4 (a)** Cumulative *prtC* abundance in saliva was compared across disease groups (Wilcoxon, p-value adjusted with BH). **(b)** Overview of replication cohorts (3,697 faecal samples in total). Two ACLD cohorts were included to validate ACLD-associated signals, two healthy cohorts and one inflammatory bowel disease (IBD) were used for specificity. **(c)** *prtC* abundance per disease group in HMP2-IBD cohort (CD = Crohn's disease, UC = ulcerative colitis, Wilcoxon). **(d)** *PrtC* gene neighbourhood in oral-gut translocators, including their phylogenetic relationship (tree on the left, based on *de-novo* assemblies). The number within each gene indicates the number of occurrences at this position relative to the center gene (*prtC* gene in orange, where the number indicates the number of assembled copies recovered from different samples). **(e)** ROC and PR curves of disease predictions (ACLD vs controls) based on prtC faecal abundance. **(f)** Faecal *prtC* abundance is significantly increased in severe cirrhosis (CLD-Sole2021, Wilcoxon). **(g)** Inoculation of human oral *Veillonella* and *Streptococcus spp.* increases intestinal barrier dysfunction in a mouse model of fibrosis. Intestinal permeability (measured by colon albumin level) compared across the non-treated group (control), $CCl_4$-treated group ($CCl_4$) and the $CCl_4$-plus gavage group ($CCl_4$+isolates) [one-sided Wilcoxon].

#### Figure S5

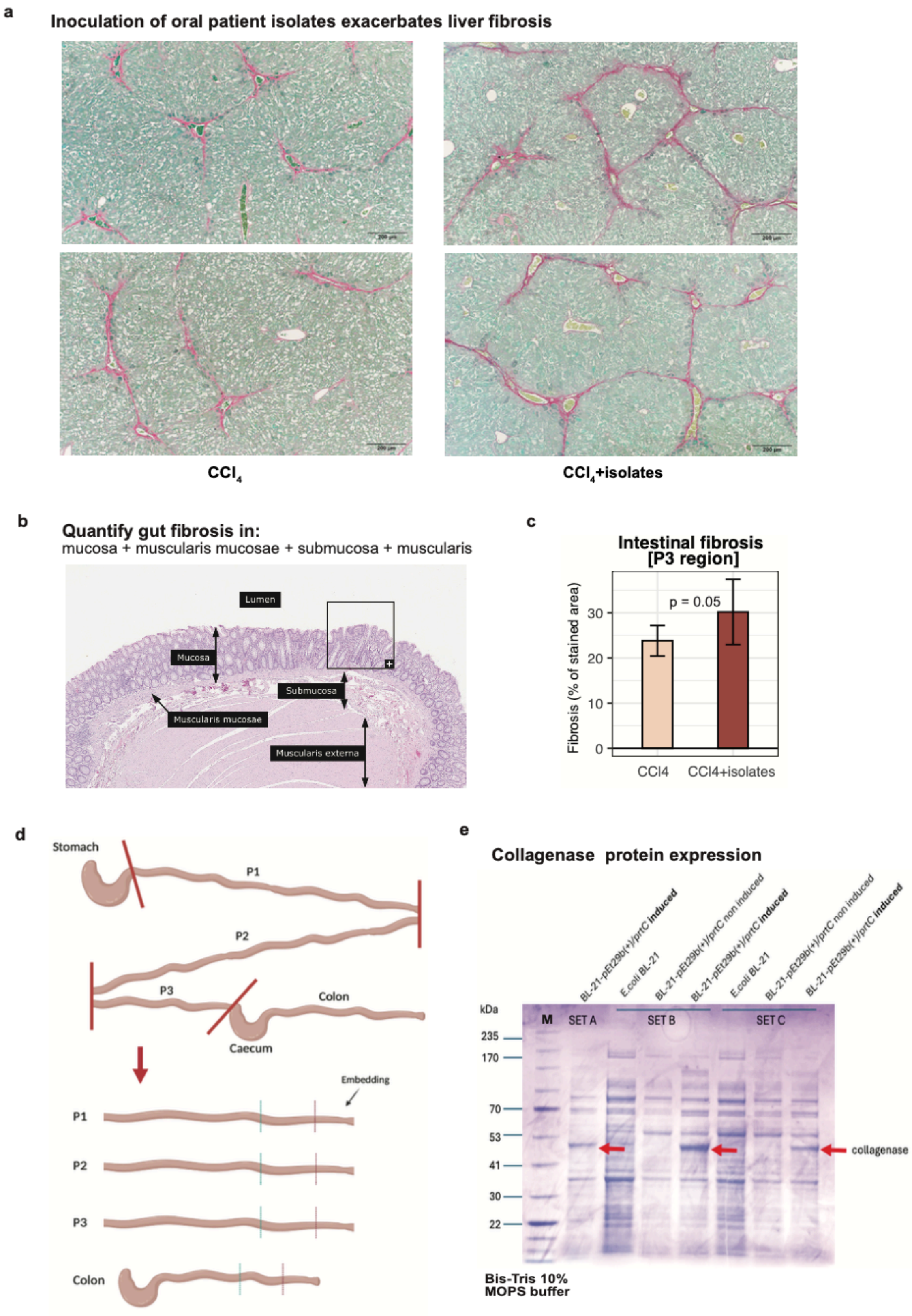
a
Inoculation of oral patient isolates exacerbates liver fibrosis
CCl4
CCl4+isolates
b
Quantify gut fibrosis in:
mucosa + muscularis mucosae + submucosa + muscularis
Lumen
Mucosa
Submucosa
Muscularis mucosae
Muscularis externa
c
Intestinal fibrosis
[P3 region]
p = 0.05
Fibrosis (% of stained area)
0
10
20
30
CCl4
CCl4+isolates
d
Stomach
P1
P2
P3
Colon
Caecum
Embedding
P1
P2
P3
Colon
e
Collagenase protein expression
BL-21-pEt29b(+)/prtC induced
E.coli BL-21
BL-21-pEt29b(+)/prtC non induced
BL-21-pEt29b(+)/prtC induced
E.coli BL-21
BL-21-pEt29b(+)/prtC non induced
BL-21-pEt29b(+)/prtC induced
kDa
M
SET A
SET B
SET C
235
170
70
53
41
30
22
collagenase
Bis-Tris 10%
MOPS buffer

**Figure S5 (a)** Inoculation of human oral *Veillonella* and *Streptococcus spp.* exacerbated hepatic fibrosis. Liver tissue was fixed, paraffin embedded and stained with Sirius red to detect fibrosis. Representative microscopic images at 10x magnification are shown, with collagen stained in red indicating regions of fibrosis. **(b)** Histological section of the gut stained with Sirius Red, showing labeled anatomical layers including lumen, mucosa, muscularis mucosae, submucosa, and muscularis extern. **(c)** Barplot showing a significantly higher percentage of Sirius Red-stained fibrotic area for the P3 region for $CCl_4$+isolates compared to the $CCl_4$ groups (p=0.05, t-test). **(d)** Schematic representation of mouse gastrointestinal tract dissection showing isolated regions: stomach, duodenum (P1), jejunum (P2), ileum (P3), caecum and colon. (**e**) Recombinant collagenase protein expression in *E.coli* BL-21 (DE3) resulted in a band of the expected size (48kDa), which was missing for the non-induced strains. M: Protein size marker; 10% SDS-PAGE stained with coomassie brilliant blue R250.

#### Supplementary Tables:

**Table S1: Bacterial strains used in this study and oral-gut translocation events details.**
**Table S2: Metadata, clinical measurements and microbial profiles**
**Table S3: Measurements from $CCl_4$ mouse experiments**

#### Reference:

Rojas-Tapias, D. F., et al. (2022). "Inflammation-associated nitrate facilitates ectopic colonization of oral bacterium Veillonella parvula in the intestine." Nature microbiology **7**(10): 1673-1685.
